# Pathogen-Phage Geomapping to Overcome Resistance

**DOI:** 10.1101/2025.10.01.679739

**Authors:** Camilla Do, Keiko C. Salazar, James D. Chang, Justin R. Clark, Austen L. Terwilliger, Paul Ruchhoeft, Paul Nicholls, Anthony W. Maresso

## Abstract

The rise of antibiotic resistance has renewed interest in bacteriophages as therapeutic alternatives. However, co-evolution of phage and bacteria will naturally give rise to phage-resistant pathogens, complicating phage therapy efforts. A critical bottleneck in the production of phage therapeutics is the discovery of virulent phages against resistant pathogens. Conventional methods for discovery are time-consuming, biased, and laborious, limiting the potential for identifying suitable phage candidates.

To overcome these limitations, we combined small-volume environmental sampling with 16S rRNA sequencing to identify reservoirs where bacterial hosts co-exist with their phage predators. This strategy, which we term geographical phage mapping (geΦmapping), pinpoints ecological “hotspots” for targeted phage hunting. We further developed a portable phage hunting device (ΦHD) that generates highly enriched phage concentrates directly from these reservoirs. By integrating geΦmapping with high-throughput enrichment, we constructed the RΦ library, a diverse collection of novel phages. We captured and isolated 36 new phages targeting extremely-resistant organisms across various ESKAPE pathogens when conventional phage hunting and experimental evolution approaches failed.

## Introduction

The rise of antibiotic resistance has brought global healthcare to the precipice of a post-antibiotic era, driving interest in bacteriophage (phage) therapy as an alternative strategy to combat resistant bacterial infections.^1–3^ Unlike antibiotics, phages evolve with their host; both locked in a relentless evolutionary arms race. Bacteria and phage continuously expand their repertoires of resistance and counter-resistance strategies. Among many examples, phages counter bacterial defenses by exposing receptors with enzymes, evading restriction enzymes with DNA base modification and evolving anti-CRISPR proteins.^4,5^ In response, bacteria resist phage infection by blocking adsorption and nucleic acid injection and using restriction enzymes and CRISPR-Cas9 to target phage nucleic acids.^6,7^ This dynamic interplay of phage resistance poses a barrier to effective phage therapy.^8^ TAILΦR is a phage center dedicated to providing lytic phages to physicians to treat patients battling multidrug-resistant infections.^9^ Across 263 distinct cases encompassing 462 bacterial isolates, 22% of isolates given to TAILOR remain untreated despite an extensive library of over 325 phages targeting 15 bacterial species (**Fig, 1A**). Information regarding TAILΦR’s phage and bacterial library can be found in **Supp. Tables 1-3**. When no phages in the library can target a pathogen, TAILΦR turns to directed evolution (to expand host range of a given phage) or environmental sampling (to find natural phages). Such approaches often fail to yield a suitable phage and so discovering new, virulent phages that can effectively target these pathogens becomes a critical and rate-limiting step **(Fig. 1B)**. As phage therapy spreads, resistant isolates will inevitably emerge under increased selection pressure, analogous to antibiotic resistance after introduction of penicillin.^10^ This puts the onus onto phage discovery where improvements are critically needed.^11^

**Figure 1:**
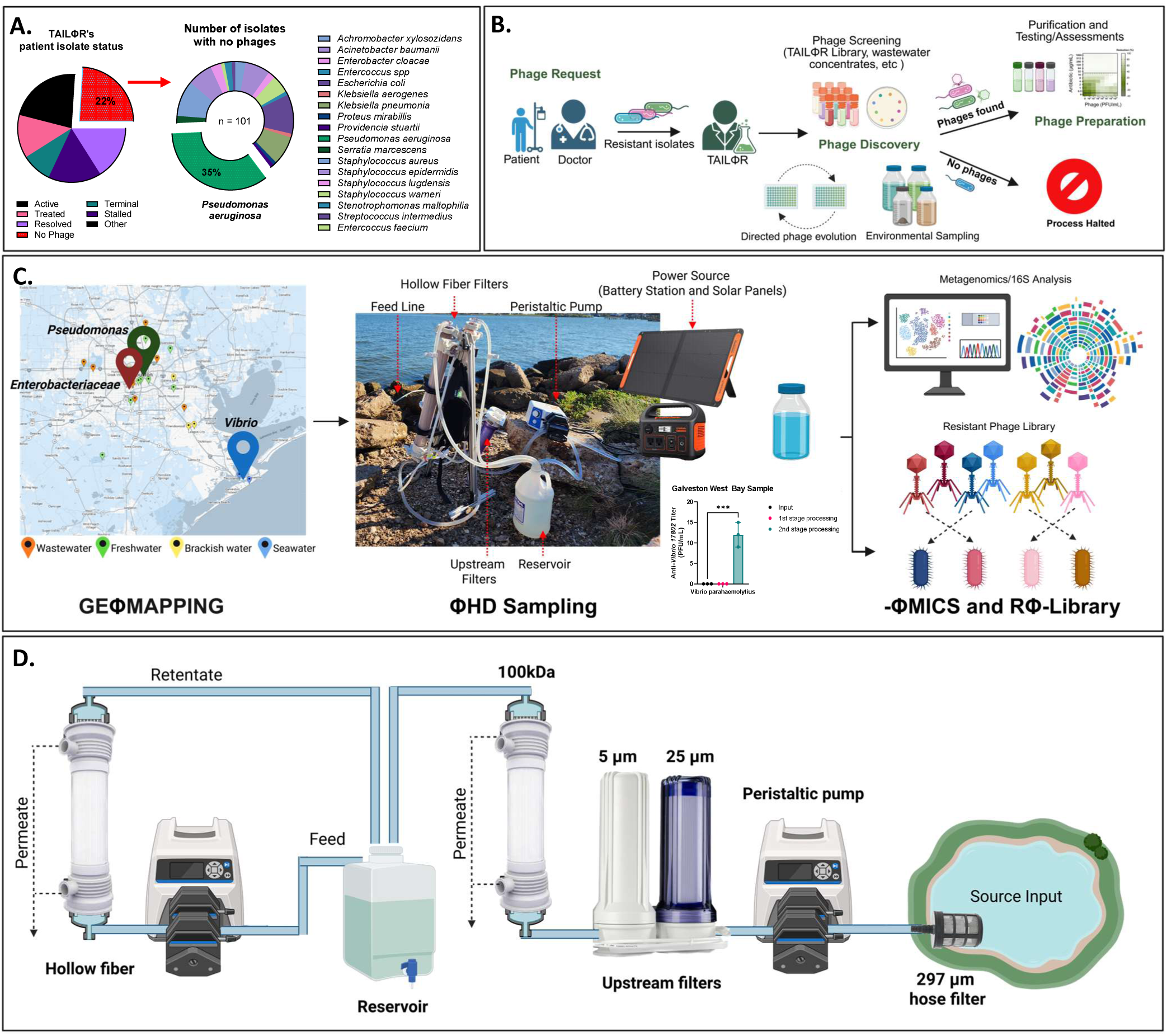
Overview and schematic of ΦHD. **A)** Pie charts of patient isolate status and percentage of bacterial strain with no phages for in TAILΦR’s library. Unfortunately, 24% of isolates that TAILΦR receives have no phages; at 35%, most of the isolates are *Pseudomonas aeruginosa*. **B**) Diagram from phage request to phage discovery and limitation. Clinicians and their patients can seek phage therapy for antibiotic-resistant infections when there are no other approved treatments available. After approval, clinicians send patient isolates to TAILΦR. TAILΦR’s phage library and wastewater concentrates are screened against the patient isolate. If phages are discovered, they are moved onto the next stage for preparation where phages are ultimately purified and tested for safety and efficacy before being handed to the clinician. When there are no phages for that isolate in the library, TAILΦR searches for phages through environmental sampling. Alternatively, they can train related phages to infect the isolate through directed phage evolution. **C)** Diagram of strategies employed in our study. Geographical phage (Φ) mapping (geΦmapping) allowed us to pinpoint target sites rich in our pathogen of interest from our environment (**left**). Once a location is chosen, we used our capture device, the phage hunting device (ΦHD), to filter, concentrate, and enrich our sample with phage and bacteria (**middle**). With the samples, we further analyzed them with phage metagenomics (ΦMICS) and screened, isolated, and characterized phage present in the samples(**right**). This led to the curation of the resistant phage (Φ) library (RΦ-Library). **D)** Schematic of ΦHD. The components on the right of the reservoir draws, filters, and concentrates water through ΦHD. On the left of the reservoir, the remaining component concentrates any material in the reservoir. Created with BioRender.com.

Phage engineering is one method of constructing phages for improved therapeutic potential. Examples include generating lytic derivatives of lysogenic phages and altering phage tail fibers to adjust their host range.^12,13^ In contrast, phage-host competition has naturally addressed phage resistance over billions of years of evolution. The challenge lies in isolating these phages from nature. Conventional methods for phage discovery can be time-consuming or introduce bias.^11^ Phage hunters are limited to sampling small volumes of water, a constraint that typically necessitates the use of enrichment techniques to increase the likelihood of phage detection.^14–17^ Enrichments can favor fast-growing or highly virulent phages, potentially overlooking those that may be therapeutically relevant but less abundant or slower to propagate.^18^ To sample larger volumes, dead-end filtration (limited by low filtration speed) and iron chloride flocculation (limited by phage inactivation) are typically used **(Supp. Table 4)**. ^19–29^ The speed and phage bias limitations are acceptable for metagenomic approaches but unhelpful for phage therapy, leaving a critical gap for patients that urgently need phages for phage-resistant bacterial infections.

Viral metagenomics involves identification of largely uncultivated viral genomes from the environment. Unfortunately, analysis is hindered by the vast diversity of viruses, lack of consensus sequences, and limited reference genomes available.^30^ Some 70% of all viral genomes are unclassified, often referred to as “viral dark matter.”^31^ Studying unknown viral genomes can advance viral ecology and may contribute useful gene products for research and therapeutics.^32^ Viral metagenomics are frequently limited by low biomass, sampling biases, and limitations in sampling remote sites.^33,34^

In this study, we sought to improve current methods for phage discovery to tackle phage-resistant pathogens and enhance gene discovery. We used a bioprospecting approach, collecting samples in East Texas, USA, and applied 16S rRNA analysis to identify pathogenic reservoirs and hence their phages. We dubbed this process, “Geographical Phage (Φ) Mapping,” or “GeΦmapping.” After identifying pathogen-rich sources, we deployed a high-throughput phage hunting device, ΦHD, for sample processing (**Fig. 1C,D**). We used this approach of combining molecular bioprospecting and high-volume sampling to address problematic “extremely phage-resistant organisms” (XΦROs) in clinical cases.

## Results

### GEΦMAPPING of water sources from Houston and Galveston, Texas

Phages are found where their hosts thrive.^35^ In marine ecosystems, for example, phage-to-bacterium ratios often reach 10:1, reflecting close associations between bacterial density and phage populations.^36^ We identified sites of high pathogen abundance to determine where to hunt for phages. We sampled 15 waterways around Houston and Galveston, Texas, USA thrice and performed 16S rRNA analysis to examine the bacterial constituents (**Fig. 2A)**. We performed rarefaction analysis to estimate species richness relative to sampling effort. The logarithmic shape of the rarefaction curve indicates that most of the abundant taxa present have been detected in each sample, although sample saturation was not fully achieved (**Supp. Fig. 1A, top**). To further quantify species richness, we compared α-diversity indices (inverse Simpson, Shannon, and Chao) and noticed that White Oak Bayou was the richest (**Supp. Fig. 1B**). To compare the membership of each site’s microbiome, we analyzed β-diversity using principal coordinate analysis (PCoA) based on Bray-Curtis dissimilarity (**Fig. 2B,C**). Samples of seawater, sewage, and freshwater clustered distinctively while samples from brackish water mingled between freshwater and seawater sites (**Fig. 2B,C**). We performed an analysis of molecular variance (AMOVA) to compare samples derived from brackish, sea, fresh, and sewage. AMOVA revealed significant structuring across all habitat comparisons, with among-group variance exceeding within-group variance in every case. Differentiation was strongest between fresh and sewage samples (Fₛ = 28.71, *P* < 0.001), followed by sea–sewage (Fₛ = 20.21, *P* < 0.001) and brackish–sewage (Fₛ = 15.92, *P* < 0.001), indicating pronounced divergence associated with wastewater. Comparisons involving brackish with seawater (Fₛ = 4.65–13.16, *P* ≤ 0.006) and brackish-fresh ((Fₛ = 5.96–13.16, *P* ≤ 0.001) were also significant (Fₛ = 4.65–13.16, *P* ≤ 0.006). Interestingly, a freshwater sample from Vince Bayou, where *Acinetobacter* was prominent, clustered with sewage samples suggesting its microbial profile resembled that of sewage. Of note, a contaminated industrial facility, formerly an oil processor and wastewater treatment plant, is situated upstream of this location.^37^ After heavy storms, toxic wastes have reportedly contaminated the area with reports of hundreds of dead fish.^38^ During sampling, toxic chemical contamination may have shifted the microbiome to resemble sewage-like communities, causing resilient genera like *Acinetobacter* to dominate.

**Figure 2:**
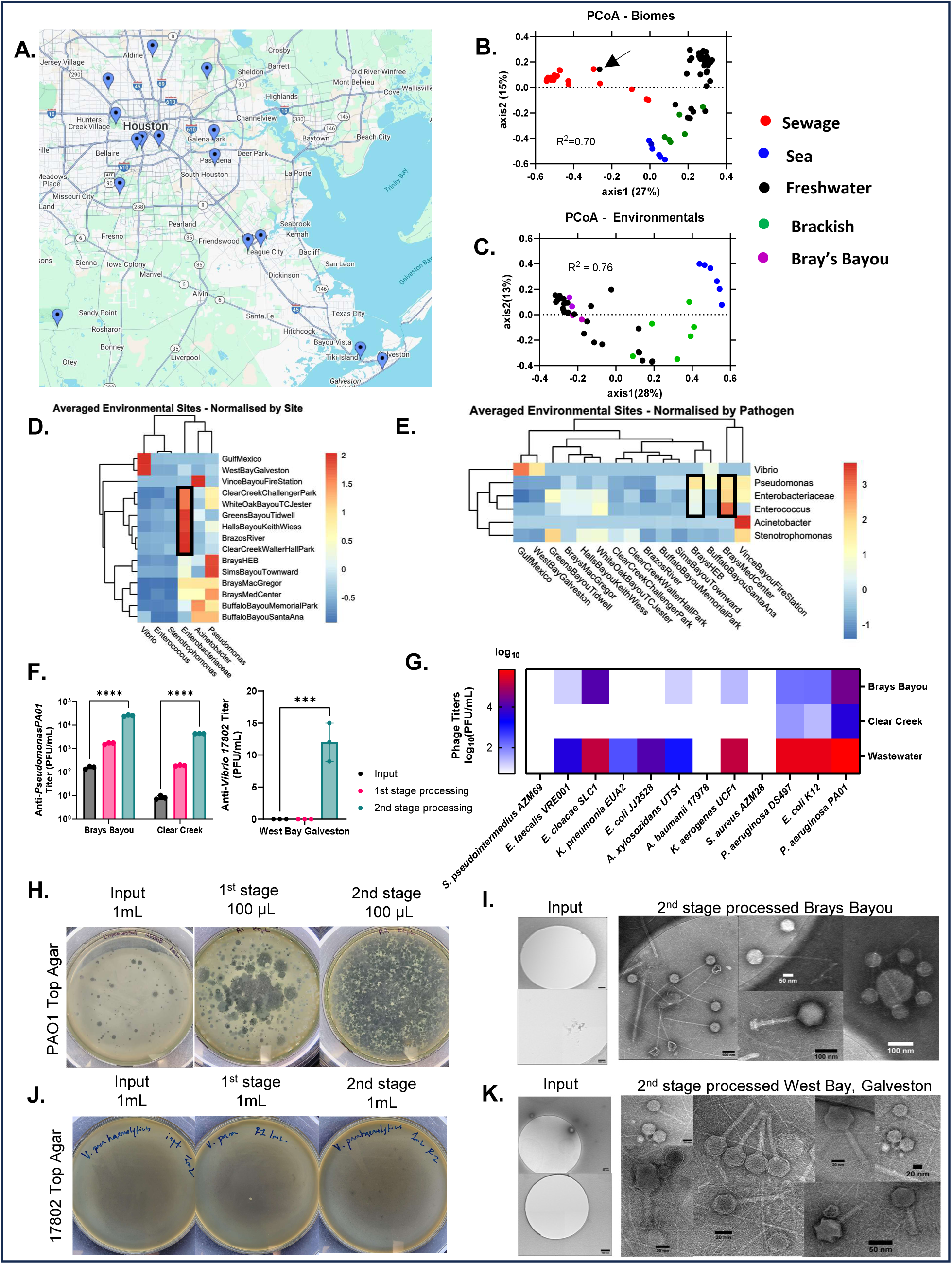
GEΦMAPPING highlights Bray’s Bayou as a Pseudomonas rich target site, and performance of ΦHD sampling of several environmental sites. **A)** Geographical map of freshwater samples obtained in Houston, TX, USA. **B)** Principal Coordinate Analysis (PCoA) analysis showing β-diversity differences between sewage, sea, and freshwater sources. The black arrow indicates the Vince Bayou sample. **C)** PCoA analysis (Bray-Curtis dissimilarity) of environmental water bodies with Bray’s Bayou target sites highlighted (purple). **D)** Heat map with z-score normalized by site shows distinct patterns of microbial prevalence at various sites. **E)** Heat map, z-score normalized by pathogen, highlights the potential of Bray’s Bayou as a target site for Pseudomonas and enteric phages. **F)** Quantification of anti-Pseudomonas (left) and anti-Vibrio (right) plaques (PFU/mL) from different stages of processing (input, first stage, and second stage processing). **G)** Heat-map showing anti-pathogen phages (log_10_(PFU/mL)) from freshwater sites. We display the mean and SEM of 3 technical repeat measurements. **H)** Plaque assay on *P. aeruginosa* PAO1 of unprocessed, first stage concentrate, and second stage concentrate of a freshwater sample. **I)** TEM of unprocessed water from freshwater. **J)** Plaque assay on *V. parahaemolyticus* 17802 of unprocessed, first stage concentrate, and second stage concentrate seawater sample. K) TEM of unprocessed water from seawater, respectively. Representative images shown (H-K).

To focus on pathogens, we extracted the counts of pathogen-related taxa and generated heatmaps normalized by site **(Fig. 2D**) and by taxa **(Fig. 2E)**. By highlighting selected pathogenic taxa, we can compare patterns across the environments relative to the other taxa; this does not imply absolute abundance or general enrichment of these taxa in the environment. Genus-level classification includes both pathogen and non-pathogenic species. Normalizing by site, we determined the microbial composition of each location. We detected *Vibrio* in Gulf of Mexico and West Bay; both expected in marine environments **(Fig 2D)**. We noted *Enterobacteriaceae* was most common in waterways draining rural and suburban areas, while *Pseudomonas* was more common in larger urban waterways (Brays and Buffalo Bayous) (**Fig. 2D, black rectangles**). Again, Vince Bayou was an outlier, highly enriched with *Acinetobacter*.

Normalizing by taxa allowed us to find sites of interest for ΦHD sampling (**Fig. 2E, black rectangles**). The data showed predominance of *Vibrio* at marine sites, while *Pseudomonas*, *Enterobacteriaceae* and *Enterococcus* were best represented at Brays Bayou. Vince Bayou remained enriched with *Acinetobacter*. Of the 22% of TAILΦR isolates without a phage, 35% of these were *Pseudomonas* (**Fig 1A, Supp. Table 3**). This made Brays Bayou, rich in *Pseudomonas,* an ideal target for high volume sampling.

### Environmental sampling freshwater and seawater with ΦHD

Although phages are abundant in nature, harvesting phages against pathogens can be difficult.^14,17^ Typically, phage hunters are restricted to small volumes and sometimes rely on enrichment techniques to compensate.^14,17^ These techniques vary by sample and host but take days with no guaranteed success. After selecting our sampling location, the next challenge was obtaining a highly concentrated sample through high-volume sampling to ensure sufficient phage abundance for effective downstream screening.

We approached this using ΦHD, which was inspired by the various research groups that used tangential flow filtration (TFF) for viral metagenomics.^25,26,39^ We designed ΦHD to have a robust filtration train combined with a high-flux concentration component, allowing for low shear fluid path and high portability (**Fig. 1D**). We used a 297 μm hose filter to screen out larger debris, followed by sequential 25 and 5 µm filter cartridges to remove larger microorganisms such as plankton and protists. We then used two hollow fiber TFF units, operating in parallel for concentration with flow rates in the low shear region. The first unit performs ultrafiltration and fills the reservoir with retentate, while the second takes from that reservoir and continuously concentrates the retentate in a closed loop. These are powered by a dual-headed peristaltic pump equipped with a lithium-ion battery and solar panels to allow prolonged operation in the field. During spike-in studies, we spiked pond water (60L) with phage (1×10^3^ PFU/mL of ΦJB10) (**Supp. Fig. 2A**). We found minimal losses of phage and bacteria during passage through the filtration train and obtained an average 20-fold increase in phage yield (**Supp. Fig. 2B**). After the spike-in study, we evaluated our cleaning-in-place protocol (**Supp. Fig. 2C, Post CIP**) for the TFF units and found it effectively removed phages and bacteria, confirming ΦHD is reusable.

In the field, we routinely achieved flow rates of 200L/hr and gathered samples ranging from 400-1200L (**Supp. Table 5**). Initial testing on-site proved the filtration/concentration train was robust enough to manage the highly polluted water ways of Houston (**Supp. Fig. 2D-E, Santa Ana Capture Site (SACS)**). Although not all *Pseudomonas* phages will be detected, we selected *Pseudomonas aeruginosa* PA01 as the indicator strain because of its broad susceptibility profile and its role as a permissive host. After first stage processing with ΦHD, phage yield increased by 95-fold in comparison to unprocessed sample (**Supp. Fig. 2E, right panel**). We also sampled the more pristine waters of Pedernales Falls (PF) and Hamilton Pool Preserve (HP), and incorporated a secondary in-lab concentration stage to further increase retentate concentrations (**Supp. Fig. 2F**). Phage yields from inputs from PF and HP were near zero but increased to 10^2–3^ PFU/mL with secondary processing. Similar treatment of SACS retentates yielded an average 3,501-fold increase in phage yield (**Supp. Fig. 2G**). This shows ΦHD, with second stage concentration, greatly increased the detection of viable phages from a given site (**Supp. Fig. 2G, 2H**). Screening with other bacteria showed more phages against other pathogens from polluted SACS water compared to pristine sites (**Supp. Fig. 2I**).

To test the combination of geΦmapping and ΦHD, we selected one site that was highlighted by our map as *Pseudomonas-*, *Enterobacteriaceae-* and *Enterococcus*-rich (Brays Bayou, **Fig. 2E**) and one low-richness site (Clear Creek). We processed 400L/site and tracked phage yields by endogenous anti-*Pseudomonas* (PAO1) phages. ΦHD significantly increased phage yields at each site (166-fold at Brays Bayou and 520-fold at Clear Creek, **Fig. 2F,H)**. Using second stage concentrates, we screened a panel of patient isolates of different genera to compare hits and found more lytic phages at Brays Bayou (7/12 species) than Clear Creek (3/12 species), phenotypically validating our 16S-based mapping strategy (**Fig. 2G**). We employed an identical filtration train at Galveston Bay sampling the marine, saltwater biome. We tracked concentrations with *Vibrio parahaemolyticus* 17802 and could only identify plaques after second stage concentration (**Fig 2F-right panel, J**), showing utility in salt water. To test whether phage concentration increased without host-selection bias, we used transmission electron miscopy (TEM). We observed a wide range of phage morphologies with intact tails in concentrates, while the input was mostly clear of particles (**Fig 2I,K**). This provides further objective evidence of unbiased concentration of intact phages through the ΦHD process.

### GEΦMAPPING of wastewater treatment plants in Houston, Texas

Pathogens belonging to *Enterococcus* and *Enterobacteriaceae,* including *Escherichia* and *Klebsiella,* are known inhabitants of wastewater, so we reasoned wastewater would be prime choice for finding phages against these pathogens.^40,41^ In a similar fashion to freshwater mapping, we sampled influents of seven different wastewater treatment plants (WWTPs) twice to generate geΦmaps of the pathogenic landscape of wastewater plants. **Fig. 3A** shows the catchment area for each WWTP in Houston. The logarithmic shape of rarefaction curves showed we adequately sampled from each site (**Supp. Fig. 1A, bottom**). PCoA of β-diversity (Bray-Curtis dissimilarity) demonstrated close clustering of sewage samples despite substantial diversity in both human populations and localization of industrial and commercial facilities. Unexpectedly, influent samples collected from Almeda Sims WWTP showed unusual abundance of *Chloroflexi*, known to be important in biodegradation in wastewater bioreactors (**Fig. 3B, yellow**).^42^

**Figure 3:**
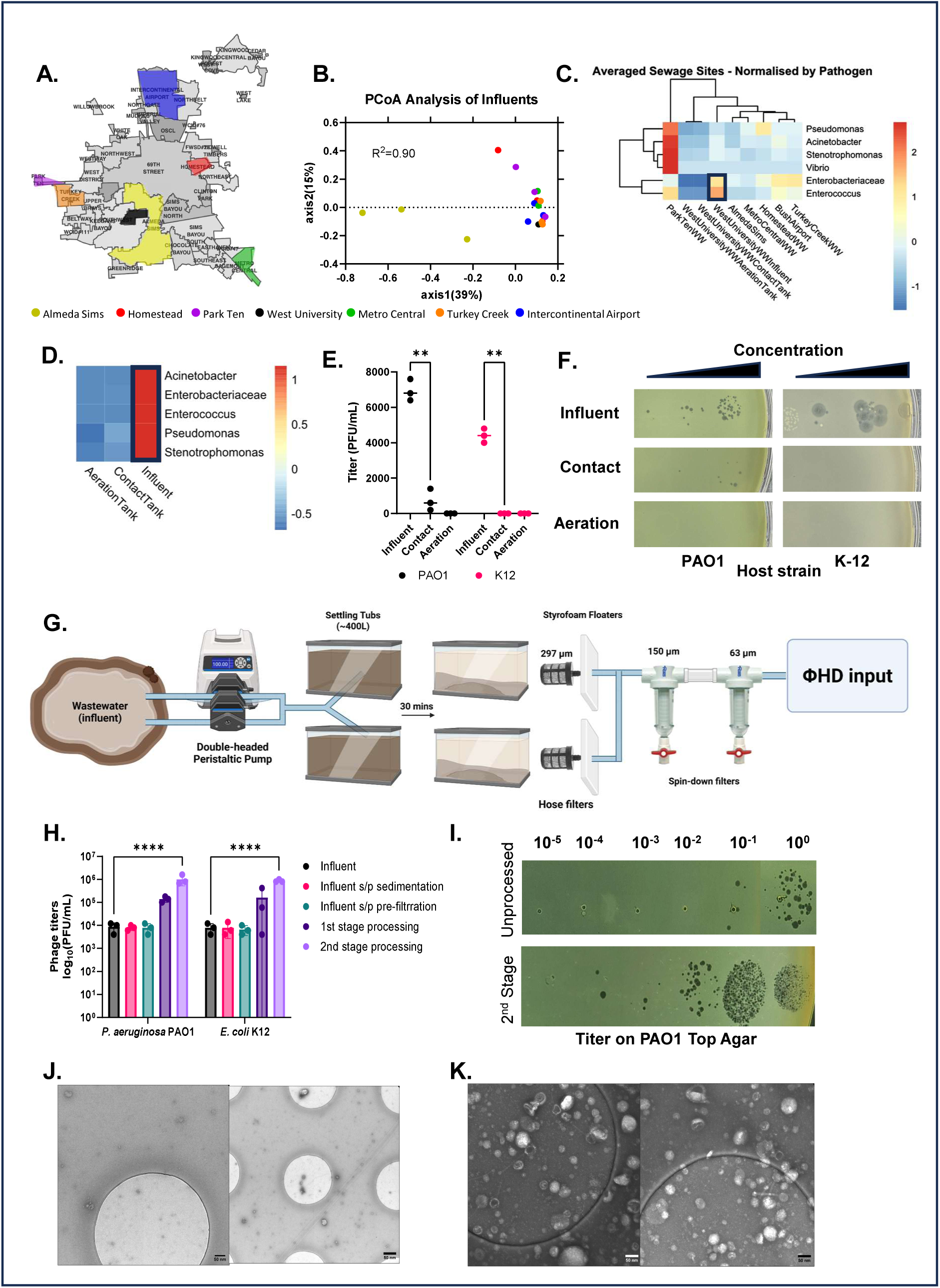
GEΦMAPPING identified *Enterococcus* and *Enterobacteriaceae* was rich at West University WWTP, and performance of ΦHD Sampling in wastewater. **A)** Schematic of geographical areas drained by WWTPs in Houston, TX, USA. **B)** β-diversity analysis shows variation in OTUs present in WWTPs by PCoA analysis based on Bray-Curtis dissimilarity (**AB,** colors represent differing WWTPs). **C)** Heatmap analysis of pathogenic taxa highlights (black box) location of enteric pathogens (z-score normalized by taxa). **D)** 16S analysis further localizes pathogenic taxa to wastewater influent (z-score normalized by taxa). **E)** Quantitation of plaques on index strains (n=3, ** = p-value ≤0.005 by 2-way ANOVA with Tukey’s multiple comparisons correction) confirms 16S sequencing data. **F)** Titration of VLPs from different WWTP processing areas on PAO1 (*P. aeruginosa*) and K-12 *(E. coli*) index strains (representative data shown). **G)** Outline of prefiltration and sedimentation step for wastewater sites. **H)** Quantification of anti-Pseudomonas plaques and anti-Escherichia plaques from different stages of processing show phage retention through the system and concentration of the end products. **I)** Titration of VLPs from different WWTP processing areas on PAO1 (*P. aeruginosa*). **J)** TEM of unprocessed ΦHD input from waastewater. **K)** TEM of second stage concentrated wastewater. Representative images shown (F, I-K).

We normalized by taxa and generated heatmaps to make our wastewater geΦmap (**Fig. 3C**). The Park Ten WWTP influent was rich in *Pseudomonas*, *Acinetobacter* and *Stenotrophomonas*, but it was being decommissioned during our study. West University WWTP had higher levels of *Enterobacteriaceae* and *Enterococcus*, so we selected it for further investigation. The wastewater treatment process alters the microbiome of wastewater, so we compared the pathogenic taxa throughout the process. We sampled raw wastewater filtered only through large steel bars that remove large debris (influent). Similarly, we sampled wastewater mixed with activated sludge (contact tank) and wastewater exposed to activated sludge with air bubbling (aeration tank).^43^ α-diversity indices showed that microbial diversity was richest in the contact tank and aeration tank of West University WWTP (**Supp. Fig. 1B**) but the influent tank had the highest abundance of the selected pathogenic taxa (**Fig. 3D**). We confirmed this 16S genomic data by comparing titers of phages against indicator strains (PAO1 for *Pseudomonas* and K-12 for *E. coli*) which showed the majority of these phages were found in the influent (**Fig. 3E-F**).

### Wastewater sampling with ΦHD

We used a modified ΦHD method to process 400L of influent wastewater from West University WWTP (**Fig. 3G**). To avoid pump issues caused by gravity, we lifted the wastewater 1–2 meters onto an elevated deck into large tubs. We allowed this material to settle for 30 minutes to sediment large debris and then we pre-filtered the sample using 297 µm hose filters and two spin-down sediment traps (150 and 63 µm screens) to produce a clarified product. This pre-filtered material was then used as the ΦHD input. To ensure these additional filtration steps did not capture significant quantities of phage, we titered the material at each stage against indicator strains (PAO1 and K12) and found negligible losses (**Fig 3H**). We titered input and second stage concentrates (**Fig. 3H,I**) on *Pseudomonas* and *Escherichia* indicators with final phage yields increasing by 118-fold and 108-fold, respectively. We attempted to look for non-selective concentration of virus-like particles (VLPs) with TEM, however this was complicated by residual particulate matter. Nonetheless, we were able to appreciate some phages in retentates but not the unconcentrated input material (**Fig 3J,K**). We also screened the second stage concentrate against the previous panel of clinical and laboratory strains. This yielded phages for 9 of 12 bacterial strains screened, and phage titers were higher than at other sites (darker red) (**Fig. 2G**).

### Collective shallow shotgun metagenomic sequencing

Phage diversity varies significantly across different biomes due to factors such as host availability, ecological niches, and environmental conditions. To test the utility of ΦHD for metagenomic applications, we compared viral metagenomes extracted from an unprocessed sample (5L) and a ΦHD concentrate (60mL of 6667X retentate, corresponding to 400L of processed sample). The majority of α-diversity indices were similar, suggesting comparable within-sample diversity; however, β-diversity based on Bray–Curtis dissimilarity (0.6062) indicates highly moderate differences in viral community composition. (**Supp. Table 6**). Heatmap of viral population abundances showed no clear clustering or distinct patterns between the two samples (**Supp. Fig. 3A**). Nevertheless, we detected substantial increases in unique viral contigs (**Supp. Table 7, Supp. Fig. 3B,C**). We compared putative *Pseudomonas*-infecting viruses from the input and ΦHD samples, and the ΦHD sample contained both different members from genetically distinct groups and a greater absolute number of unique anti-*Pseudomonas* viruses (**Supp. Fig. 3D,E,F**).

Given the apparent utility of ΦHD in concentrating samples for metagenomics, we looked at the viromes of fresh (Brays), brackish (Clear Creek), waste (West University WWTP), and seawater (Galveston). Based on α-diversity indices, either freshwater or wastewater samples represent the richest sources of viral diversity **(Supp. Table 8).** Heatmap of viral abundance and β-diversity analysis with PCoA confirm distinct viral community compositions between biomes.(**Supp. Fig. 4**). We clustered our putative, non-redundant viral sequences into vOTUs, or virus operational taxonomic units. We generated a gene-sharing network with vCONTact2 with each node representing a vOTU, colored by source and clustered them with reference genomes from the Prokaryotic Viral RefSeq v201 database (**Fig. 4A**). We counted 16,198 vOTUs ≥ 5kb and 5095 vOTUs ≥ 10kb.

**Figure 4.**
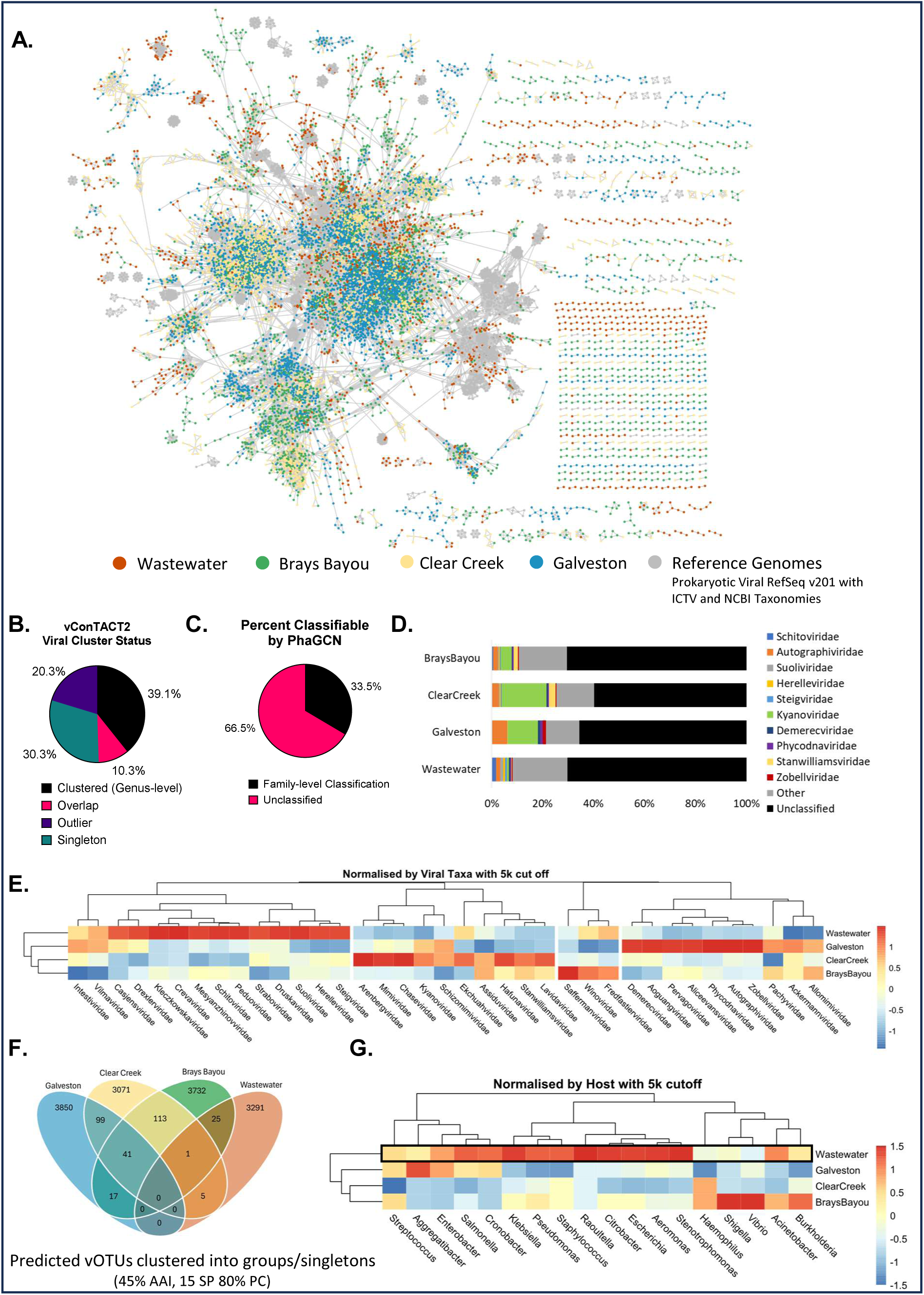
Taxonomic classification of viral contigs from metagenomic datasets reveal distinct viral signatures as well as biased presence of viruses of pathogenic hosts between different locations. **A)** vConTACT2 gene-sharing network of 16,198 viral operational taxonomic unit (vOTUs) (≥ 5kb) from West University WWTP, Brays Bayou, Clear Creek, and Galveston. Each dot (node) represents a vOTU, and each line (edge) represents the similarity between each genome. Additionally, 3,508 reference genomes from the Prokaryotic Viral RefSeq v201 database are shown in grey. **B)** Pie-chart of viral cluster status of vOTUs from vCONTact2. **C)** Pie-chart of classified (family-level) vs unclassified vOTU from PhaGCN. **D)** Bar graph of family-level classification (PhaGCN counts) of topmost abundant genera across the samples. Remaining genera are grouped together as “others” in grey. Unclassified vOTUs are in black. **E)** Heat map with z-score normalized by viral taxa to show distinct viral signatures between sites (pheatmap separated each cluster). **F)** Venn-diagram of shared vOTUs (cluster mode: amino-acid identity (AAI) 45%, protein coverage (PC) 80%) depict number of unique vOTUs present at each site and shared between sites. **G)** Heat map with z-score normalized by host reveal the relevance of sampling specific sites for phage hunting.

39.1% of vOTUs clustered with a reference genome at the genus level, while 20.3% were outliers sharing only 1-2 genes with other genomes. 30.3% were singletons with no genes related to any other genomes in the data set (**Fig. 4A,B**). We also used PhaGCN to classify each virus, but only 33.5% of viruses could be classified to the family-level (**Fig. 4C**). Of the classified genomes, *Kyanoviridae* (green) and *Autographiviridae* (orange) made up the largest identifiable taxa in Clear Creek and Galveston (**Fig 4D**). This agrees closely with previous findings that *Kyanoviridae*, containing T4-like viruses that infect cyanobacteria, and *Autographiviridae,* T7-like viruses, have been found in deep-sea viromes.^44^ Due to current limitations in viromics, a large portion of our metagenomic dataset remain unknown. Collectively, between 60.9% (outlier/singles from vCONTact2) and 66.5% (unclassified from PhaGCN) of vOTUs are in this category. A substantial fraction is considered viral dark matter, but we used the remaining identifiable sequences to provide some biological context. This enable us to validate our sampling strategies and compare samples and biomes from another. We used counts from PhaGCN classifications to generate heatmaps normalized by viral taxa to see distinct viral clusters from each biome type. There are four viral signatures present, separating wastewater, seawater, freshwater and brackish water (**Fig. 4E**). We further clustered viruses from each biome together to identify overlaps and visualized them with Venn diagrams (**Fig. 4F**). We saw that overlaps of viral genomes reflected the physical environment; for example, the brackish water of Clear Creek overlapped between seawater and freshwater - representing its status as an admixture of the two. Next, we used CHERRY for host prediction and made heatmaps normalized by hosts. A large majority of phages with pathogenic hosts were found in wastewater virome (**Fig. 4G, black box**).

To increase phage diversity and minimize the emergence of phage resistance, curating a large, genetically diverse phage library is advantageous. Such resource allows researchers and clinicians to select phages tailored to their patient needs, including targeting specific phage receptors, maximizing host coverage, designing phage cocktails, or reducing dependence on any single phage.^45^. We compared *Pseudomonas*- and *Escherichia*-infecting viruses from freshwater and wastewater. Phylogenetic analysis of *Pseudomonas*-infecting viruses showed significant increases in diversity when combining vOTUs from the two biomes (**Fig. 5A**). Counting and clustering the unique genomes, there were no sequences that were in common between sewage and freshwater (**Fig. 5B**). Analyzing the *Escherichia*-infecting viruses, phylogenetic diversity was greatly increased by including freshwater phages, and the two biomes had no sequences in common (**Fig. 5C,D**). This shows the benefit of creating a phage library using ΦHD from multiple biomes.

**Figure 5.**
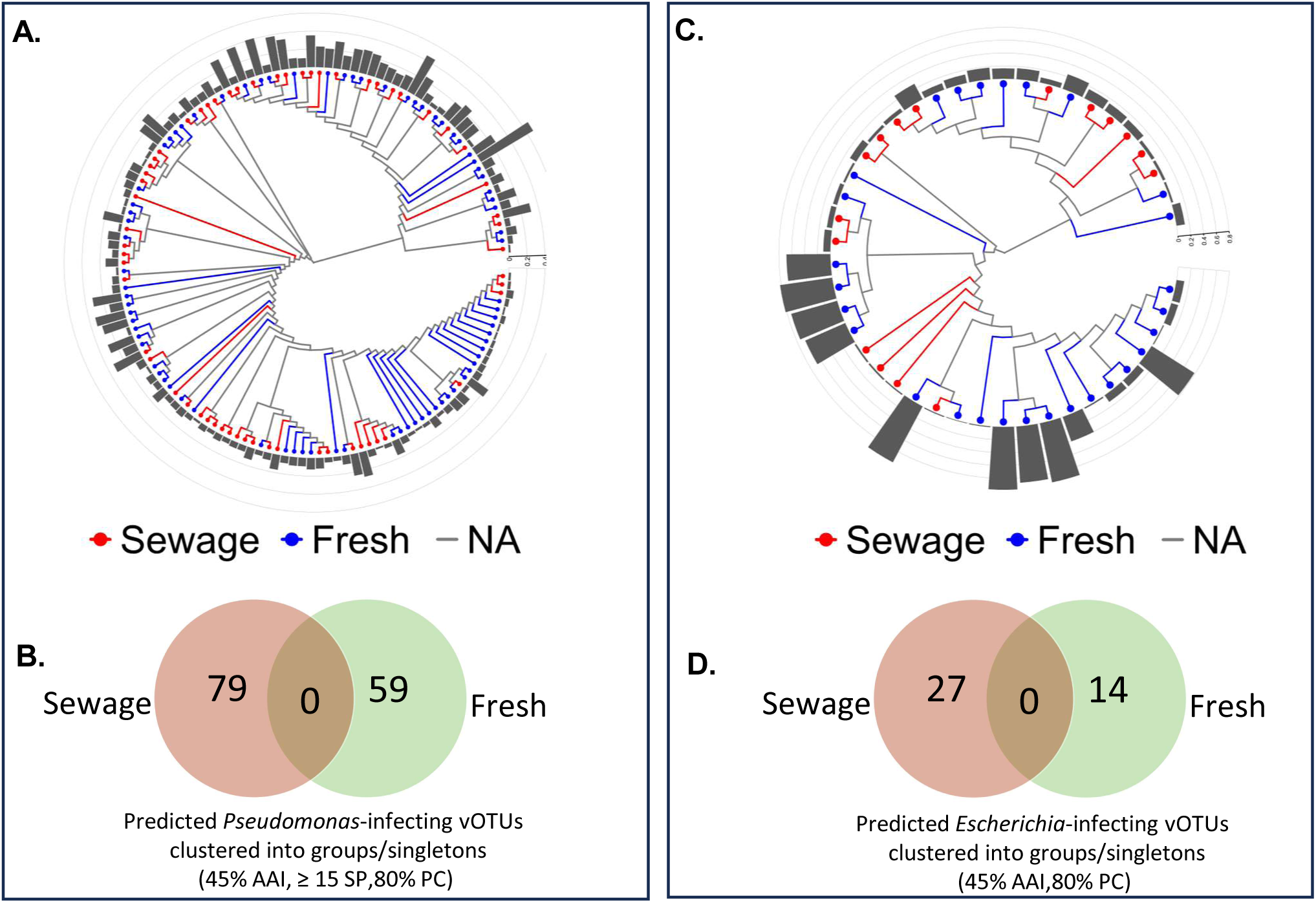
Relevance of multi-site sampling for increased diversity and collection of unique *Pseudomonas and Escherichia* phages. **A)** Phylogenetic tree of all vOTUs predicted to have *Pseudomonas* host from West University WWTP wastewater and Brays Bayou freshwater metagenomic dataset. Branch-length is non-scaled. **B)** Venn-diagram of vOTUs-grouping (cluster mode: AAI 45%, 15 shared protein (SP), PC 80%) show no common vOTUs shared between sites. **C)** Phylogenetic tree of all vOTUs predicted to have *Escherichia* host from wastewater and Brays Bayou metagenomic dataset. Branch-length is non-scaled. **D)** Venn-diagram of vOTUs-grouping (cluster mode: AAI 45%, 15 shared protein (SP), PC 80%) show no common vOTUs shared between sites. Bar graphs (A, B) represents the genomic similarities between a contig and its closest neighbor based on genome-wide sequence similarities computed by tBLASTx from VipTree.

### Resistant phage library (RΦ-Library)

Last-line antibiotics serve as a critical safety net for treating life-threatening infections by multi-drug-resistant bacteria. Their use is restricted to preserve efficacy and slow the development of drug resistance. In a similar manner, the RΦ-library was conceptualized as a last-line phage library to combat extremely phage resistance organisms (XΦROs). To test our approach of geΦmapping and ΦHD and generate a therapeutic phage library for XΦROs that posed clinical challenges, we used 17 XΦROs from the TAILΦR collection. These failed all conventional phage discovery methods (**Fig. 6A**) including sewage spotting and Appelmans protocol (for host range expansion). As screening ΦHD concentrates allows searching far larger volumes (**Fig. 6B**), we immediately found therapeutic phage candidates for all 17 XΦROs using freshwater (Brays Bayou) and sewage (West University) sources (**Fig 6C**). We purified (**Fig. 6D**) and characterized 36 new phages by genomic analysis and TEM (**Fig. 6D,E, Supp. Table 9).** This collection of phages is what we termed the RΦ-library. Compared to TAILOR’s sequenced phages, proteomic dendrograms (**Fig. 6E**) showed phages with both subtle and high genetic variation, including new genera exemplified by MYC30C1 and MYC30C2 against *Enterococcus* (**Supp. Fig. 5, 6**). Our technique also isolated the jumbo phage UCS29C1 against *E. coli*, which was closely related to the *Klebsiella* jumbo phage Miami, showing that even large phages can survive the process intact (**Supp. Fig. 5, 6**).^46^ These successes make GEΦMAPPING and ΦHD sampling excellent additions to TAILΦR’s phage discovery pipeline, allowing for isolation of novel phages with greater precision and efficiency (**Fig. 6F**).

**Figure 6:**
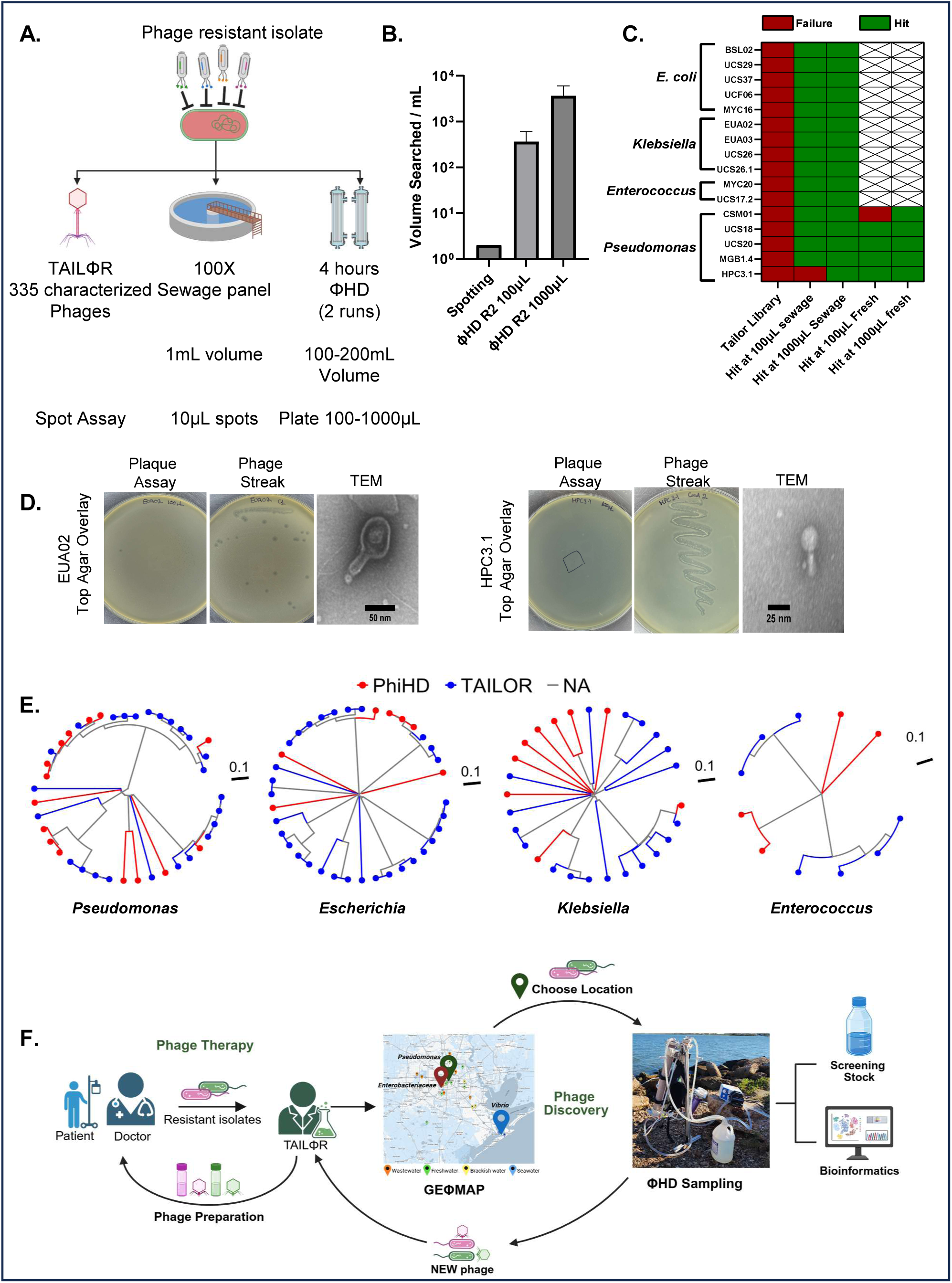
ΦHD increases search volumes and allowed efficient discovery of phages for phage-resistant bacterial pathogens. **A)** Schematic of ΦHD compared to standard identification and discovery of therapeutic phages, **B)** Effective volumes that can be searched comparing standard spotting of concentrates with high-concentration, high volume ΦHD retentates (n=2 retentates with differing final concentrations), **C)** Heat map showing success/failure of ΦHD compared to the TAILΦR library in finding therapeutic phage candidates. **D)** Plaque assays (**left column**) on phage-resistant strains EUA02 (*K. pneumoniae*) and HPC3.1 (*P. aeruginosa*). Individual plaques (middle columns) were streaked and isolated from plaque assays. TEM images (right columns) of select isolated phages infecting EUA02 and HPC3.1. Representative images shown. **E)** Dendrograms of all isolated phages for *Pseudomonas*, *Escherichia*, *Klebsiella*, *Enterococcus*. Phages from ΦHD are in red, and phages from the TAILΦR library in blue. **F)** Diagram of the implementation of ΦHD and geΦmapping to the phage discovery pipeline. Created with BioRender.com.

## Discussion

Our study leverages the billion year-long evolutionary war between phages and their hosts, focusing on discovering and harnessing natural phages. We report: i) the design, construction, and successful use of a high-throughput phage capture device (ΦHD); ii) the use of this device to identify regional sources of both bacteria and phages (geΦmapping); iii) the use of geΦmapping coupled with ΦHD sampling to identify novel phages that target phage-resistant strains, when all other phage matching solutions failed (RΦ-Library); and iv) the characterization of new metagenomes. Collectively, this work provides a rich resource for therapeutic phages against resistant ESKAPEs (***E****nterococcus faecium, **S**taphylococcus aureus, **K**lebsiella pneumoniae,* ***A****cinetobacter **b**aumannii, **P**seudomonas aeruginosa, and **E**nterobacter spp.)* and a rich gene library for new biological discovery (**Fig 6F**).

Wastewater, areas of high human activity and dense urban development are particularly rich in phages capable of targeting antibiotic-resistant pathogens.^14,16,47–49^ This is largely due to the elevated microbial loads in these settings, which naturally support a higher abundance and diversity of phages.^50,51^ Human activity imposes evolutionary pressure on bacteria, driving the emergence of antibiotic-resistant strains—and, by extension, their phage counterparts.^52^ To strategically identify optimal sampling locations, we employed geΦmapping to pinpoint environments with high pathogen loads (**Fig 2E, 3C**). We identified Brays Bayou as an excellent locale for anti-*Pseudomonas* and other phages, with agreement between genomic mapping and phenotypic phage surveys. We extended our approach to sewage, not only localizing to a single WWTP for phage hunting but to a specific section of that plant (**Fig. 3 A-F**).

This targeting enabled the use of ΦHD which produced phage-rich retentates with phage yields increasing by 100-3500-fold each sample in comparison to unprocessed input (**Fig. 2F, 3H, Supp. Fig. 2G**). These retentates were used to target XΦROs in clinical cases resulting in not only the generation of the RΦ-library (**Fig. 6C**) and discovery new genera of phages (**Supp. Fig. 5, 6**) but also treatment in patient cases. Currently, 1 patient has received 2 phages that necessitated the ΦHD approach, and 6 more patient cases are in progress that required ΦHD-derived phages (not published). This illustrates the critical need of phage discovery being addressed by the geΦmapping and ΦHD approach.

Our study is limited by multiple factors. Our current geΦmap is limited to the Gulf Coast area around Houston, TX, USA and thus is of limited utility to others. However, the overall approach of localizing phage reservoirs is generalizable. With regards to the geomap-guided phage hunt, we only compared one low host-availability control site (Clear Creek) with two high host-availability sites (wastewater and Brays Bayou). More comparisons, especially incorporating even more sites with varied host availability and including additional hosts to screen, would strengthen the claim. Beyond our control, we were limited by weather conditions, as temperature and rainfall can drastically alter both microbial and viral composition. In turn, this limited our sampling time to days with consistent weather (i.e., dry, warm days) for accurate comparisons. In the field, it is technically challenging to remove all phages and bacteria from the hollow fiber units at the end of a sampling run. We employ simple backflushing through the retentate port which increases phage yield by 25-50%. More phages could be stuck in the hollow fibers. Given the amount of particulate removed later during cleaning, we are certain recovery percentages can be further increased with additional backflushing. While the ΦHD system is portable, the total assembly is large and requires at least two people to carry and operate. We are currently developing a smaller, portable version for single person use to address this. This would allow for researchers to sample in more remote locations and require less manpower. Finally, our approach is limited to aquatic biomes, but it is well known that the soils of forests and other areas (animal waste, sand, and so on) harbor huge phage biodiversity. We are currently working on ways to address this for both viral ecology and clinical phage discovery.

Collectively, we used a bioprospecting approach to localize therapeutic phages and harvest them *en masse* with ΦHD. This resulted in large metagenomic data sets informing the viral ecology of the Gulf Coast, critical therapeutic phages for challenging clinical cases and tools for others to use in phage discovery.

## Methods*

### Sample acquisition and preparation for 16S mapping

4L freshwater and seawater samples from 15 locations were collected in containers cleaned with 10% bleach and copiously pre-rinsed in ddH_2_O. Locations and weather were precisely recorded (**Supp.Table 10**). Maps were made with EasyMapMaker.^1^ Samples were collected in triplicates with the exception to Brays Bayou @ HEB. Based on 16S mapping and data not shown here, this location is biologically equivalent to Brays Bayou @ Medical Center due to its short distance and lack of inflow between them. For sampling purposes, we used this location due to its easier access to the bayou and safety reasons. Samples were transported back to the laboratory within hours of gathering samples and stored at 4°C until further processing next day. We centrifuged approximately 400 mL of each sample at 10,000xg for 20 minutes to pellet bacteria and debris. 250 mg was measured out in pre-weighed tubes. In some samples, where there were insufficient pellets (e.g. Buffalo @ Santa Ana Capture Site, Galveston, West Bay, Sims Bayou, Halls @ Keith Weiss Park), we centrifuged an additional 400mL of water until 250 mg was met. Sewage samples were obtained from the influent tank of various wastewater treatment plants by plant workers and transported to BCM and handled in a similar fashion. We centrifuged 50 mL wastewater influent at 10,000xg for 20 minutes. 250 mg of the pellet was measured in pre-weighed tubes.

### DNA preparation and 16S processing

DNA was extracted using the DNeasy PowerSoil Kit (Qiagen, Germany) according to the manufacturer’s instructions. Quality was evaluated by A260/280 ratio using a NanoDrop ND-1000 spectrophotometer (Thermo Fisher, USA). Samples with a A260/280 ratio outside of the 1.8-2.0 range were further processed using a Monarch Genomic DNA Purification kit (NEB, USA) using the “clean-up” protocol in the manufacturer’s instructions. Samples were reprocessed from the start if DNA concentrations were insufficient. Samples were sent for sequencing to Novogene. The V4 region of the 16S gene was amplified using the 515F (5’-GTGCCAGCMGCCGCGGTAA-3’) and 806R (5’-GGACTACHVGGGTWTCTAAT-3’) primers^2,3^, and then sequenced on a Novaseq6000 and 100K raw reads collected (Novogene, China). We included negative controls with each sequencing batch, consisting of the PowerSoil kit without sample to generate a “kitome” to assess the laboratory and kit background. For these control samples, 16S rRNA amplification and sequencing was completed despite lack of high-quality DNA inputs. Raw sequences were uploaded to SRA (BioProject #: PRJNA1309115).

### 16S data analysis

Raw reads were processed in mothur (v1.48.0) guided by the MiSeq SOP using SILVA (v132) reference files and seed.^4–7^ Owing to the size, number of samples, and relative lack of computational resources we used the cluster.split command with taxlevel=4 to cluster the sequences into OTUs. We generated rarefaction curves using default parameters and calculated alpha-diversity statistics in mothur, subsampling 50,000 random sequences from each group. Beta diversity analysis and PCoA was likewise performed in mothur using Bray-Curtis distance matrices and visualized in Graphpad Prism (v.10.1.2, GraphPad Software, USA). Using R, we performed AMOVA to confirm our β-diversity comparisons. To analyze and visualize the identified taxa in each sample we generated Krona charts on a Galaxy web server.^8,9^ We generated heatmaps of the relative amounts of specified taxa by calculating z-scores in R.^10^ We calculated z-scores both by site and by taxa, and displayed both using pheatmap.^11^

### ΦHD environmental sampling

A filtration/concentration device, ΦHD, was designed to filter microbes and phages from water. The filtration train starts with a 297 µm stainless steel mesh hose filter (3.6’’ in height and 2’’ in width) attached to silicone peristaltic tubing and goes into a water source (the feed line). The hose filter excludes trash, plant matter, and fish from the system. The tube connects to a Countertop Filtration System equipped sequentially with a 25 µm blown polypropylene cartridge and a 5 µm spun polypropylene cartridges. The filters remove more debris (soil, sand, etc.) and larger microorganisms like bacteria. Tubing then connects to the feed port of a 100 kDa hollow fiber unit (modified polyether sulfone (mPES), 65 cm, KrosFlo Hollow Fiber Filter Module). Anything smaller than 100 kDa (viruses, smaller bacteria, etc.) is filtered by tangential flow filtration (TFF) and filtered through the permeate ports. The retentate port ejects filtered water into a (pre-cleaned with 10% bleach and diH_2_0-rinsed) 4L reservoir. Stop valves are connected to the retentate line and bottom permeate line to control flow rates and fill the jacket of the unit. We use one head of a dual-headed peristaltic pump to propel water through the filtration train. The pump is powered with a travel-size battery station, Jackery Explorer 300, that was charged with solar panels.

Water was filtered and concentrated into a single reservoir, and a second 100 kDa hollow fiber unit was used to further concentrate the water into smaller volumes. The second head of the pump continuously cycled retentates from reservoir to unit. We used a custom-built aluminum stand that allows easy field use of the hollow fiber units. The concentrated product was expected to be 500 mL to 1 L in volume. ΦHD sampling in-field is the first stage of processing (**Supp. Fig 7A**). Samples were brought back to the lab in 1-2 hours. We centrifuged the samples, generally 1-2L, at 10,000xg for 20 minutes and filtered the samples with 0.22 µm vacuum-filters (Steritop filters (PES). Samples were stored at 4°C until second stage processing in lab (next day). A smaller version of the 100 kDa hollow fiber unit was used to achieve volumes of 50-200 mL per sample (mPES, 20 cm, MidiKros Hollow Fiber Filter Module). Retentate flow rates are controlled by Luer lock stoppers on the retentate line and bottom permeate line. Concentrations of each sample were recorded accordingly after secondary concentration (**Supp. Table 5**).

### Cleaning-in-place (CIP) protocol of hollow fiber units (TFF units)

After each ΦHD run, we cleaned the TFF units to regenerate them. For 5-10 mins, water was flushed through the permeate ports on both ends of the TFF unit. This reverse flow removes the majority of biological build-up in the units. The outer jacket and hollow fibers were then filled with 0.5M HCl and left overnight to incubate at room temperature. The next day, we rinsed the units with water then soaked outer jacket and hollow fibers in 0.5M NaOH overnight. Once rinsed with diH_2_O, the units are ready to be used again. For more polluted waters (such as wastewater), we repeated the CIP protocol. We’ve successfully used the units over 10 times for this study (**Supp. Table 5**).

### ΦHD wastewater sampling

Wastewater has a high burden of flotsam (toilet paper, wet wipes and feminine hygiene production, etc.) as well as smaller particulates (**Supp. Fig. 7B**). We pre-filtered wastewater and used the pre-filtered input as the feed to ΦHD. Two silicone peristaltic tubes drew wastewater 1-2 meters from the influent tank of West University WWTP and into two large tubs, pre-cleaned with 70% ethanol and Clorox wipes. This helped avoid gravitation issues such as slower flow rates and battery tripping. Wastewater was allowed to settle for 30 minutes. This allowed larger, denser material to sediment to the bottom. Two tubes with styrofoam floaters and barbed hose filters propelled water through two spin down filters – containing 150 and 63 µm mesh cartridges. The floaters were essential in avoiding the settled debris at the bottom, and the larger filters helped remove more debris. Wastewater was stored in two pre-cleaned tubs; this became the input for ΦHD where we processed the water as described above. For wastewater processing, an additional valve on the feed line is needed to slow down flow rate. Similarly, we centrifuged the first stage concentrate (1-2L) at 10,000g for 20mins and filtered the samples with 0.22 µm vacuum filters. second stage processing lowered the volume to 150-200 mL.

### Bacterial growth and plaque assays

A table of strains used in the study can be found in **Supp. Table 11**. From a single colony, bacterial cultures were inoculated in Luria Broth (LB, Sigma-Aldrich) and grown overnight at 37°C with shaking at 180 rpm. For plaque assays, overnight cultures were suspended in 0.75% LB top agar with 10-1000µL of ΦHD concentrate and poured onto 1.5% LB plates to solidify. Plates were incubated overnight at 37°C. Plaques are counted next day. Phage titers were visualized with Graphpad Prism (v.10.6.1, GraphPad Software, USA). For phage streaks, a single plaque on agar plates was picked and resuspended in 25 µL of phosphate buffer saline. Bacterial cultures were suspended on 0.75% LB top agar and poured onto LB plates. Before the top agar solidifies, the resuspended plaque was streaked with a pipette. To isolate individual phages from a plaque, the phage was continuously streaked out until plaque morphologies are consistent (at least twice).

### DNA preparation and shotgun shallow metagenomic sequencing

11.5 mL of second stage concentrates from ΦHD (**Supp. Table 5)** were centrifuged with a SW-41Ti rotor at 37.5k rpm for 2 hours to pellet virus-like particles (VLPs). Supernatant was removed, and the pelleted viruses are used for DNA extraction using the DNeasy PowerSoil kit (Qiagen, Germany) according to manufacturer’s protocols. The extracted DNA was sent for PCR-free library preparation followed by shallow shotgun metagenomic sequencing on the NovaSeq X Plus system platform using the PE150 strategy (Novogene, Beijing). This resulted in >2G of raw data per sample with Q30>85. Raw sequences were uploaded to SRA (BioProject #: PRJNA1308632).

### Viral metagenomic analysis

Raw data was uploaded into KBASE^12^ and examined with FastQC (0.12.1)^13^ then trimmed with Trimmomatic (v0.36)^14^ with default settings (sliding window size 4, sliding minimum quality 15, post tail crop length null, head crop length 0, leading minimum quality 3, trailing minimum quality 3, minimum read length 36).^14^ Post-trimmed reads were assessed with FastQC (0.12.1). Reads were assembled into contigs with metaSPAdes (v3.15.3); minimum contig length was 300 ≤ 2000 bp and k-mer sizes were 21,33,55,77,99, and 127.^15^ Contigs were uploaded into CyVerse for further analysis with VIBRANT.^16,17^ VIBRANT was used to identify and extract viral contigs from the dataset; default settings were used (length of bp was 1000, number of ORFs per scaffold was 4). Viral contigs from individual samples were clustered into viral operational taxonomic unit (vOTU) using average nucleotide identity (ANI)-based clustering (95% ANI, 85% alignment coverage) on PhaBOX.^18^ We used CheckV^19^ to assess quality and completeness of vOTUs. vOTUs were annotated with PROKKA (v1.14.5)^20^ on KBASE, and gene-sharing networks were made with vConTACT2 (v0.9.19).^21^ Settings were set to default, and the reference database used for cluster taxonomy was NCBI Bacterial and Archaeal Viral RefSeq V201. We visualized the gene-networks with Cytoscape.^22^ vOTUs were further taxonomically classified with PhaGCN^23^ (minimum length ≥ 5kb, 75% amino acid identity (AAI), shared protein 15, and 80% protein coverage) and predicted host of each vOTUs with CHERRY^24^ (minimum length ≥ 5kb, 75% amino acid identity (AAI), shared protein 15, and 80% protein coverage, 90% CRISPRs identity). Again, we generated heatmaps of the relative amounts of identified taxa or predicted hosts by calculating z-scores and displayed them using pheatmap in RStudio.^10,11^

vOTUs with predicted hosts in the genera *Pseudomonas* and *Escherichia* were manually extracted from the datasets of wastewater (West University WWTP) and freshwater (Brays Bayou). We clustered vOTUs with AAI-based clustering mode on PhaBOX (45% AAI, 15 shared protein, and 80% protein coverage). We used VIPTree^25^ phylogenetic analysis then visualized the resulting Newick trees using ggtree^26^ in RStudio.

### Statistical analysis of metagenomic sequences

Raw data was uploaded into KBASE and examined with FastQC (0.12.1) then trimmed with Trimmomatic (v0.36) with default settings (sliding window size 4, sliding minimum quality 15, post tail crop length null, head crop length 0, leading minimum quality 3, trailing minimum quality 3, minimum read length 36). Post-trimmed reads were assessed with FastQC (0.12.1).

We used MetaPop to analyze viral metagenomic sequence data at the interpopulation (macrodiversity) level.^27^ We aligned and mapped filtered, trimmed reads of each biome to our reference genome with bowtie2 (v2.5.4+galaxy0). The 16,198 vOTUs (95% ANI, 85% alignment coverage) from wastewater, seawater, brackish, and freshwater were combined and used as our reference genome. Generated BAM output files from bowtie2 and the reference genome were used as input for MetaPop. MetaPop’s macrodiversity analysis includes raw population abundance, normalized population abundance, and α-diversity calculations. The following analyses were conducted in RStudio using tidyverse, vegan, and pheatmap. From normalized population abundance tables, we calculated z-scores and derived heatmaps to display feature-level patterns. We calculated β-diversity using Bray-Curtis dissimilarity, then we performed PCoA with the ordination of the Bray-Curtis matrix. To assess robustness, 2,000 features were randomly subsampled and analysis repeated across 1,000 bootstrap iterations. Resulting ordinations were aligned to a reference with Procrustes alignment. Mean coordinates and standard deviations were calculated for each sample, and scatter plots were generated.

### Generation of RΦ collection

We obtained clinical isolates from the collection of TAILΦR Labs at Baylor College of Medicine (BCM). We selected isolates for which no phage had been isolated despite exhaustive searches in sewage spotting and testing of the TAILΦR phage library. Study of these clinical isolates was approved by BCM’s Institutional Review Board (IRB). We grew each isolate overnight in LB media, mixed 100 μL of this overnight culture with 3 mL of 0.7% top agar and either 100 μL or 1mL of environmental or sewage ΦHD concentrate as indicated. Plates were incubated overnight at 37°C. Next day, plaques that we considered as lytic in appearance (non-cloudy, cleared zones on bacterial lawns) were picked with a pipette tip and streaked in top agar with the same clinical strain prepared in an identical manner. Plaques were streaked 3 times to ensure purity. We endeavored to pick 2-3 different plaque morphologies per clinical isolate for each source (environment or sewage). The purified plaques were picked and mixed in 100 μL of phage buffer (10 mM Tris pH 8.0, 100 mM NaCl, 10mM MgSO_4_). We then made plate lysates by mixing 50 μL of this phage-laden material with 100 μL bacterial overnight of the indicated strain and 3 mL 0.7% top agar and incubating overnight. These were harvested by scraping the top agar from the plates, rinsing the plate surface with 3 mL phage buffer and centrifuging this mixture at 2200xg for 10 minutes and removing residual bacteria with a 0.22 μm PVDF filter (Merck Millipore, Ireland). These phages were titered using standard methods on 0.75% top agar on the indicated strain. Anti-HPC3.1 phages underwent plate lysate production in a laboratory strain PAO1 owing to the amount of mucus produced, and were subsequently titered on the indicated clinical strain. Any other phages found in this study was noted in **Supp. Table 12**.

### Genetic analysis of the RΦ collection

We employed various methods of DNA preparation for the phages. For samples with adequate titers, we used the EZNA Universal Pathogen Kit (Omega Bio-Tek, USA) according to the manufacturer’s instructions. Phage strains with lower titers or that failed to produce adequate DNA using the EZNA kit were processed by generation of 9 mL of plate lysate (as above) and centrifugation with a Sw-41Ti rotor at 37.5k rpm for 2 hours to pellet virus-like particles (VLPs). This pellet was then processed using a Qiagen Power Soil Kit using the manufacturer’s instructions. For DNA samples with ratios outside of 1.8-2.0, we further purified them with the NEB Genomic DNA Purification kit using the manufacturer’s “clean-up” protocol. Whole genome sequencing using standard protocols was done by BCM’s CMMR core. We processed short reads data using FastQC (v0.12.1)^13^ to assess data quality, removed low quality sequences (Q>30 cut off) and removed the adapters using Trimmomatic (v0.39)^14^ with default settings. We verified removal of adapters and quality cutoffs with another run of FastQC. These clean reads were then assembled using SPAdes (v3.15.3) (DNA source standard, minimum contig length 200 ≤ 500)^28^ and annotated using RASTtk (v1.073) (default setting). We identified candidate viral sequences from these assemblies using ^29^VirSorter2 (discarding sequences with <2 hallmark genes, requiring hallmark genes on all sequences, only output high confidence viral sequences, and requiring viral genes to be annotated).^30^ Sequences with high dsDNA phage score, >4-6 hallmark genes, relatively low cellular gene fraction and coverage suggestive of high copy number above the bacterial host were examined manually and BLASTn^31^ was used to determine phage or not. Confirmed phage sequences were then re-annotated with RASTtk (v1.073) to provide final phage genomes for the RΦ library. We compared TAILOR sequences (pre-existing data) with the sequences of purified phages from RΦ library. We used VIPTree^25^ for a proteomic dendrogram and then visualized the resulting Newick trees using ggtree^26^ in RStudio. Several Klebsiella phages that were isolated on related host strains were identical and so we removed these duplicates prior to visualization. All phage genomes were re-annotated with Pharokka v 1.9.0.^32^ Coding sequences were predicted with PHANOTATE.^33^ tRNAs were predicted with rRNAscan-SE 2.0.^34^ tmRNAs were predicted with Aragorn.^35^ CRISPRs were predicted with CRT.^36^ Functional annotations were generated by matching each CDS to the PHROGs^37^, VFDB^38^, and CARD^39^ databases using MMseqs2^40–42^ and PyHMMER.^43^ Phage genomes were rotated with Dnaapler.^44^ Contigs were matched to their closest hit in the INPHARED^45^ database using mash.^46^ GenBank accession numbers for phage genomes are on **Supp. Table 9** and **Supp. Table 12**.

### Electron microscopy and figures

Various environmental, sewage and purified phage samples were imaged at BCM’s Cryo-EM Advanced Technology Core Facility. Samples were deposited on grids (Quantifoil 2/2 200 Cu + 2-nm ThinC; Quantifoil Micro Tools GmbH, Jena, Germany) and negatively stained with 2% uranyl acetate using standard techniques. Images were collected with either a JEOL 1230 electron microscope (JEOL, Japan) at 80 kV equipped with a 4,000 by 4,000 Gatan Ultrascan charge-coupled device camera (Gatan, Ametek) or a JEOL 2100 (200 kV) electron microscope equipped with a Gatan 4k x 4k CCD. Images were collected at numerous magnifications, and representative examples were selected and viewed in NIH ImageJ. We used BioRender to generate **Figure 1** and **Figure 6F**.^47,48^

## Acknowledgements

This work was supported by the National Institutes of Health (grant number 5 U19 AI157981) with additional funding from the Levy-Longenbaugh Fund and Kleberg Foundation. PN received additional support from T32 AI0554413 fellowship, K08 AI173452-01A1, and BCM Intramural funds. We thank Alleigh Nicholls, Tai Gip, Christian Maresso, Dylan Chirman, Ling Qiu, Ellen Vaughan, John Taylor and Rachel Lahowetz for their assistance on in-field sampling. CryoEM data was collected at the Baylor College of Medicine CryoEM ATC, which includes equipment purchased under support of CPRIT Core Facility Award RP190602.

