## Supplementary Figures and Tables for "Pathogen-Phage Geomapping to Overcome Resistance"

### Supplemental Figure 1.

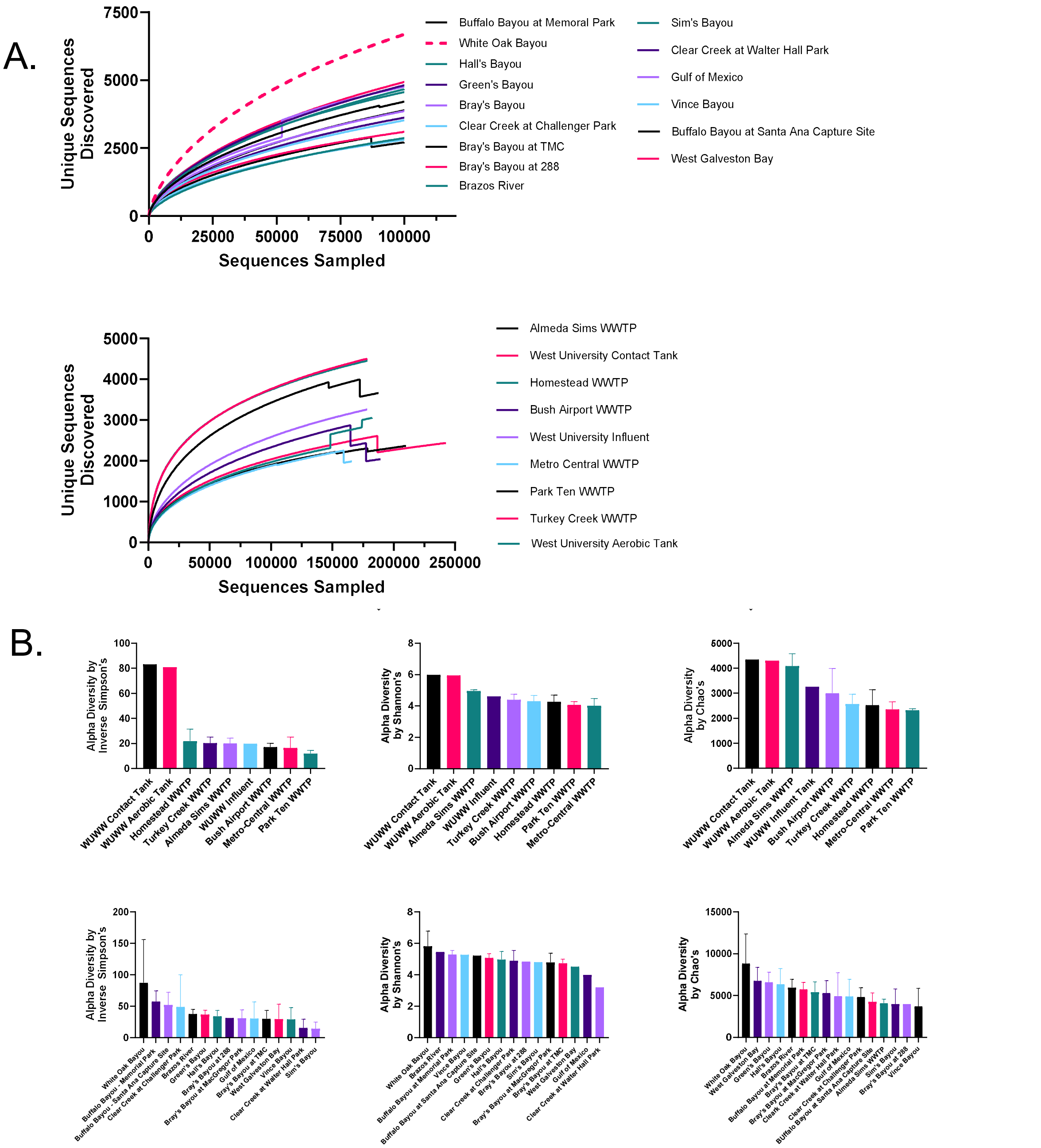


**Supplemental Figure 1: 16S analysis show adequate sampling (rarefaction curves) and high diversity (α-diversity indices) for sampling collection**. **A)** Rarefaction curves for each site show no sites were under-sampled in environmental sites **(top panel)** or sewage sites **(bottom panel)**. Curves represent the mean of triplicate biological samples for all sites except for West University WWTP. Rarefaction was performed in MOTHUR with the default 1000 randomizations. **B)** α-diversity indices for sewage sites only (**top panel**) and environmental sites only (**bottom panel**). Left panels show the Inverse Simpson's Index, middle panels show the Shannon index and right panels show the Chao's index. Averages represent biological triplicates, except for West University measurements where only single replicates were available, and error bars show SEM. α-diversity calculations were made using MOTHUR where we standardized the calculation to 50,000 random sequences for each group.

### Supplemental Figure 2.

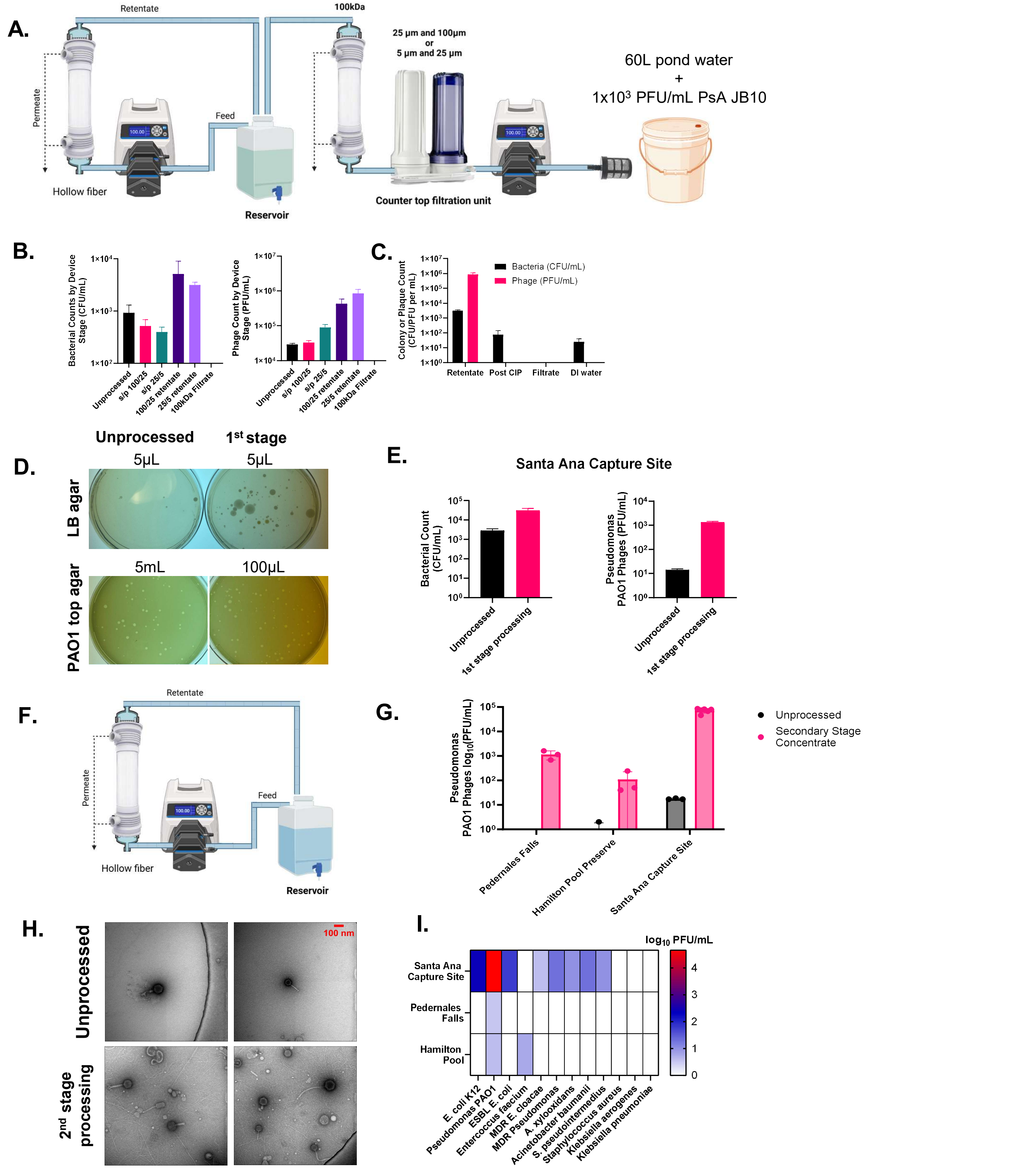


**Supplementary Figure 2: Spike-in and early on-field ΦHD sampling experiments**. **A)** Schematic of ΦHD spike-in study where 60L of pond water was spiked with 1x10$^3$ PFU/mL of *Pseudomonas* phage JB10. We used two set-ups with changes in the upstream filters: 100/25 µm or 25/5 µm set-up. **B)** Bacterial counts (**left**) and phage titers (**right**) were measured for each upstream filtration (status post s/p 100/25 or s/p 25/5) and after concentration with ΦHD (retentates). Indicator strain used was *P. aeruginosa PAO1*. **C)** Validation of cleaning in place (CIP) protocol on PAO1 after spike-in study. **Mean and SEM from 2-4 technical replicates are shown.** With ΦHD, we performed our field test at Santa Ana Capture Site (SACS). **D)** Representative plates of unprocessed and first stage concentrated samples from the Santa Ana Capture Site (SACS), plated on LB agar or Pseudomonas PAO1 lawns. **E)** Quantification of anti-Pseudomonas plaques and bacterial colonies from SACs from two separate runs. We incorporated an in-lab, benchtop concentration step to our ΦHD retentates**. F)** Schematic of second concentration phase of the ΦHD system in-lab. Using ΦHD, we sampled freshwater from Pedernales Falls (PF), Hamilton Pool (HP), and SACS. **G)** Quantification of anti-Pseudomonas phages from PF, HP, and SACS before processing and after second stage processing second **H)** TEM imaging shows sparse virus-like particles (VLPs) in unprocessed SACS samples compared to concentrated VLPs in SACS retentate samples. Representative images shown. **I)** Heat-map showing anti-pathogen phages from each sampled site. We display the mean and SEM of 3 technical repeat measurements.

**Supplementary Figure 3:**

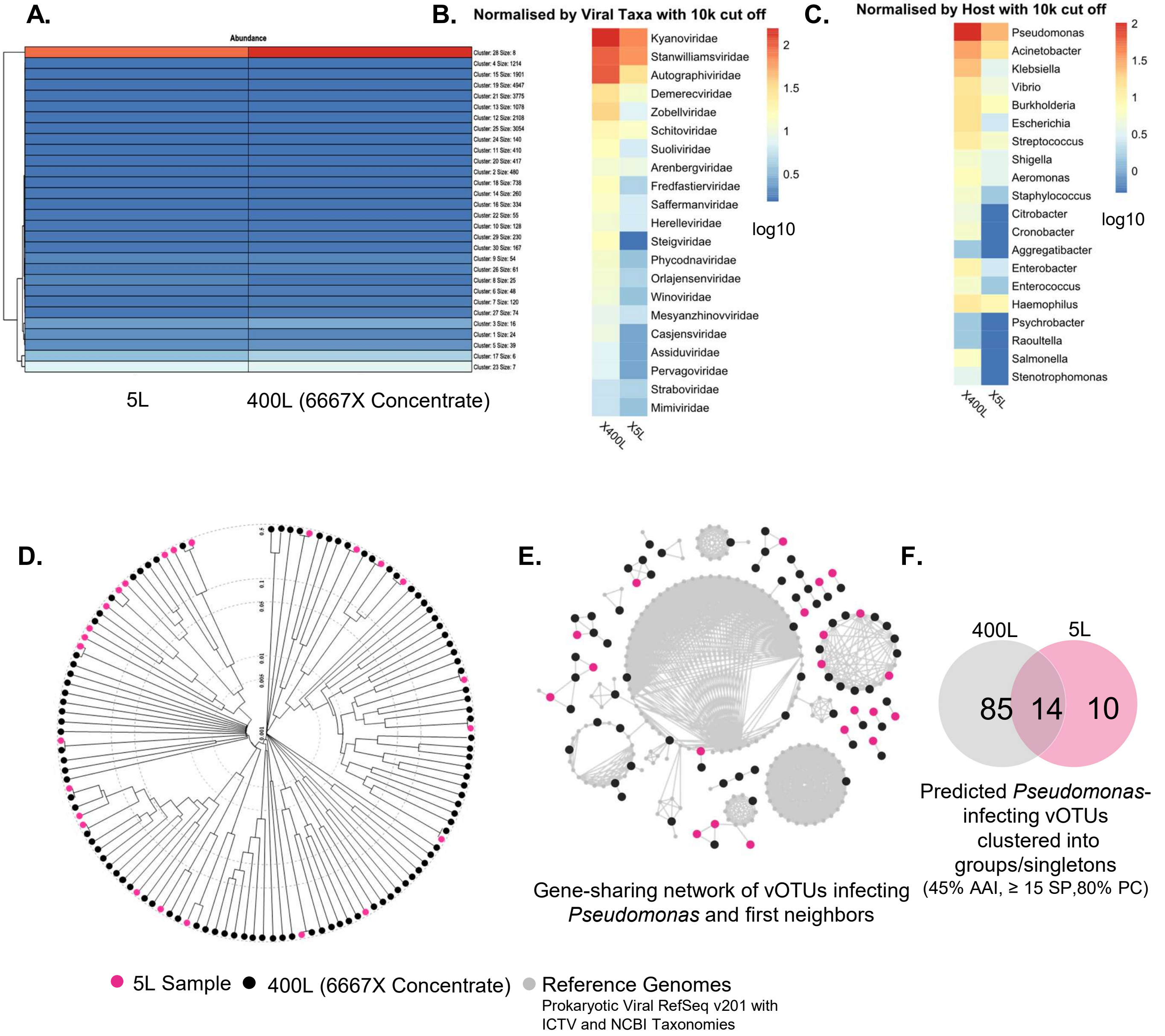


**Supplementary Figure 3: ΦHD concentrate (processed 400L) has a greater number of viruses present than unprocessed sample (5L). A)** Heatmap of viral population abundances, calculated using z-scores. The 21918 vOTUs identified from the 400L and 5L freshwater samples (Supp. Table 8) were used as the reference genomes for MetaPop's macrodiversity pipeline. **B)** Heatmap normalized by viral taxa (10kb cut-off for vOTUs). We classified vOTUs with PhaGCN to the family-level, and generated heatmaps to visualize the concentration of taxas from a 5L vs 400L sample. **C)** Heatmap normalized by predicted hosts (10kb cut-off for vOTUs). We predicted hosts of each vOTU and generated heatmaps to visualize the concentration of pathogenic taxa from each sample. **D)** Phylogenetic tree of all vOTUs predicted to target *Pseudomonas* hosts from 6667X (100 vOTUs) and 5L unprocessed sample (25 vOTUs). Greater than half of the vOTUs from the 5L sample have some relatedness to the vOTUs from the 400L ΦHD-concentrate. **E)** vConTACT2 network analysis of clustered vOTUs infecting *Pseudomonas* and first neighbors, including reference genomes. **F)** Venn-diagram of vOTUs-grouping (cluster mode: AAI 45%, 15SP, PC 80%) show shared vOTUs between samples.

**Supplementary Figure 4:**

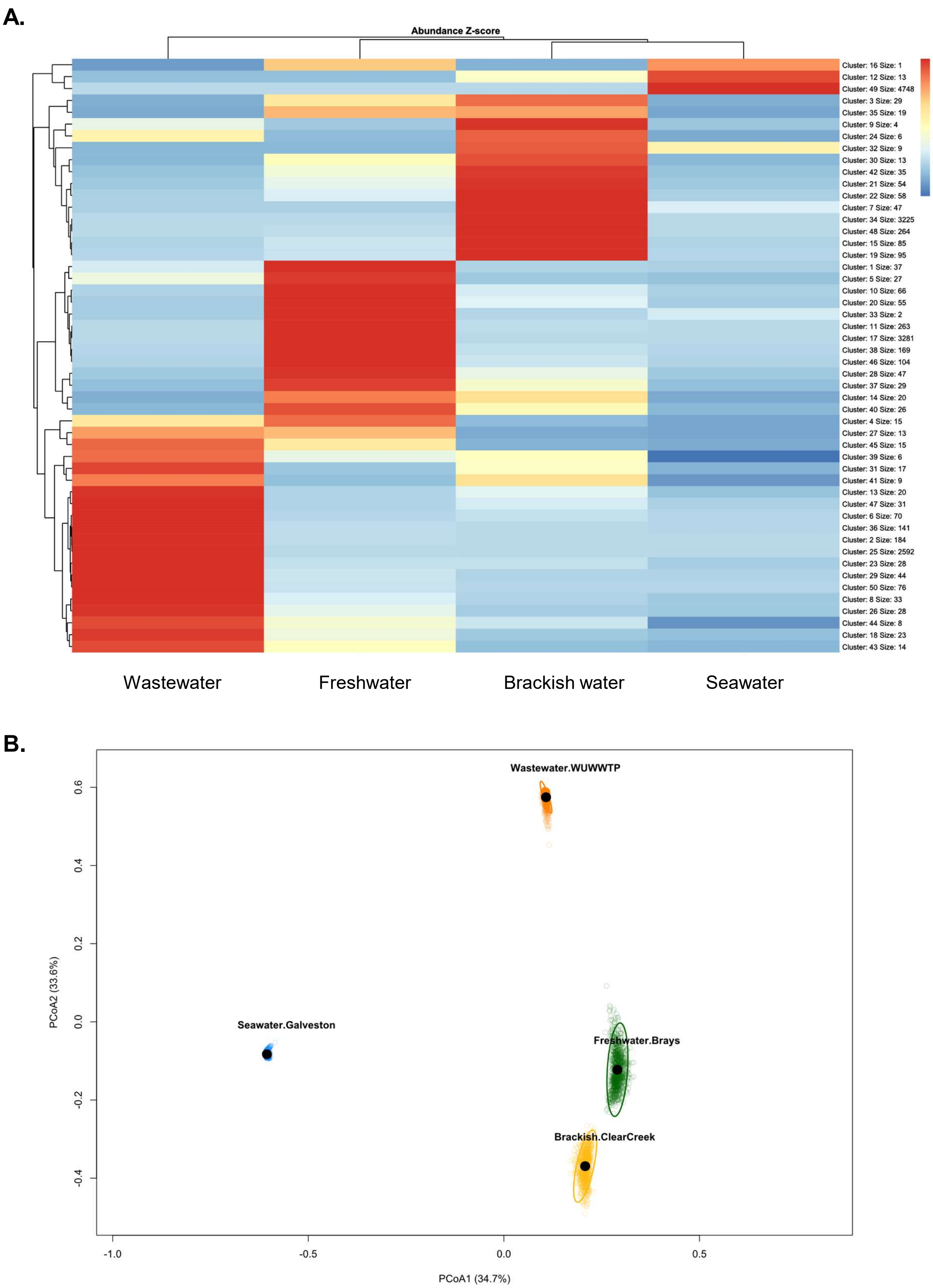


**Supplemental Figure 4: Viral community patterns across biomes. A)** Heatmap of z-scored viral population abundances reveal distinct feature-level patterns between biomes. Abundance tables were generated using the MetaPop macrodiversity pipeline with reference genome set as the collection of all vOTUs identified across the four biomes. **B)** PCoA (based on Bray-Curtis dissimilarity) compares viral communities in seawater (blue), freshwater (green), brackish water (yellow), and wastewater (orange). For PCoA, 2,000 features were randomly subsampled and analysis repeated across 1,000 bootstrap iterations. Resulting ordinations were aligned to a reference with Procrustes alignment. Mean coordinates and standard deviations were calculated for each sample.

**Supplemental Figure 5.**

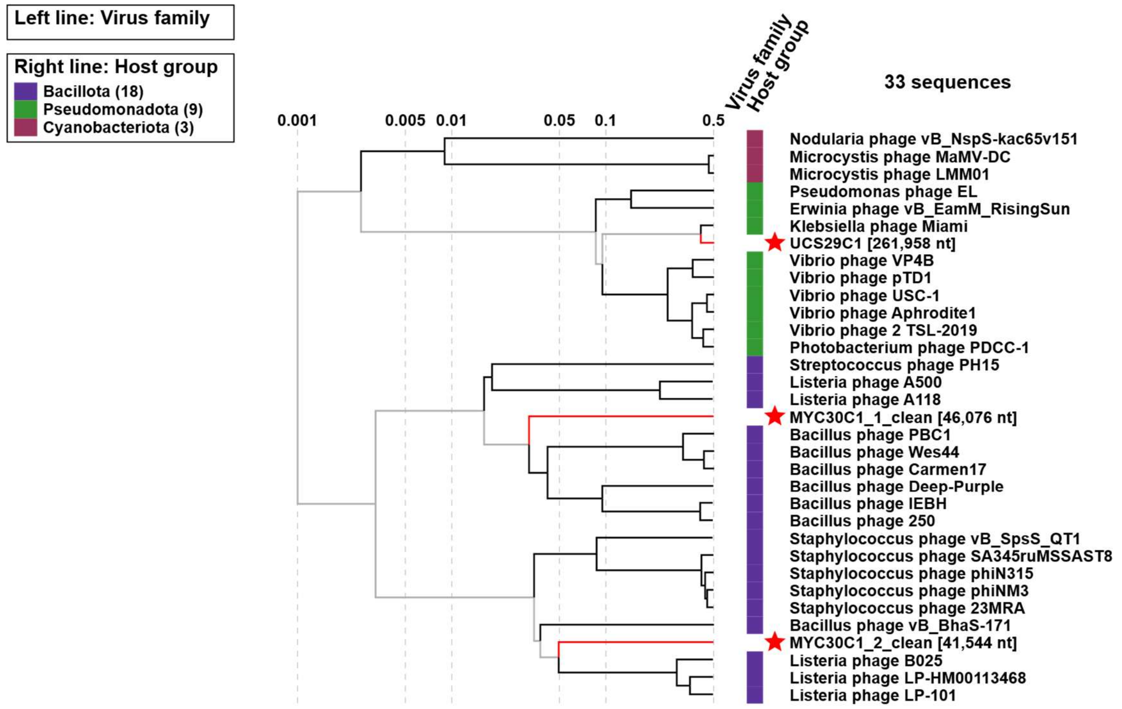


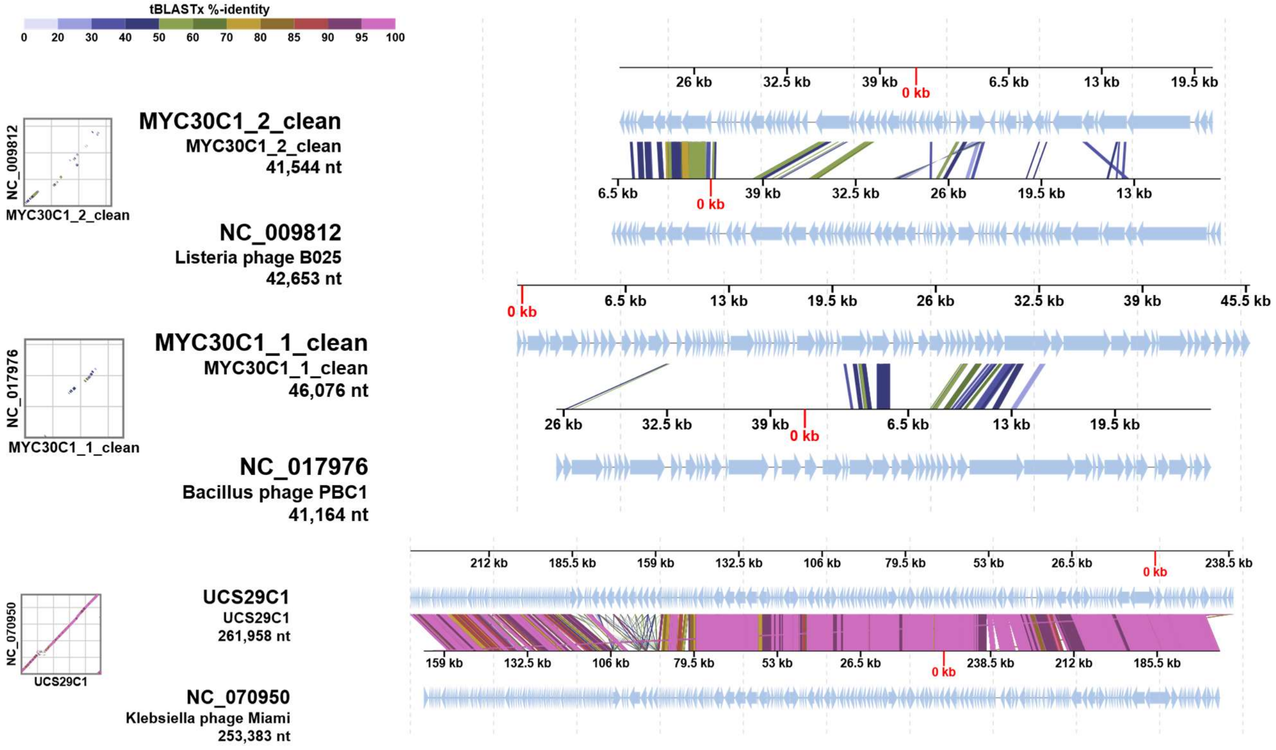


**Supplementary Figure 5: Unique phages isolated from GEΦMAPPING and ΦHD sampling.** Proteomic clustering (VIPTree) of UCS29C1, MYC30C1_1 and MYC30C1_2 with their closest neighbor found through BLASTn to look at how distinct their genomes are (**top panel**). Aligned phages with their closest neighbor to depict their percent identity (**bottom panel**).

**Supplemental Figure 6.**

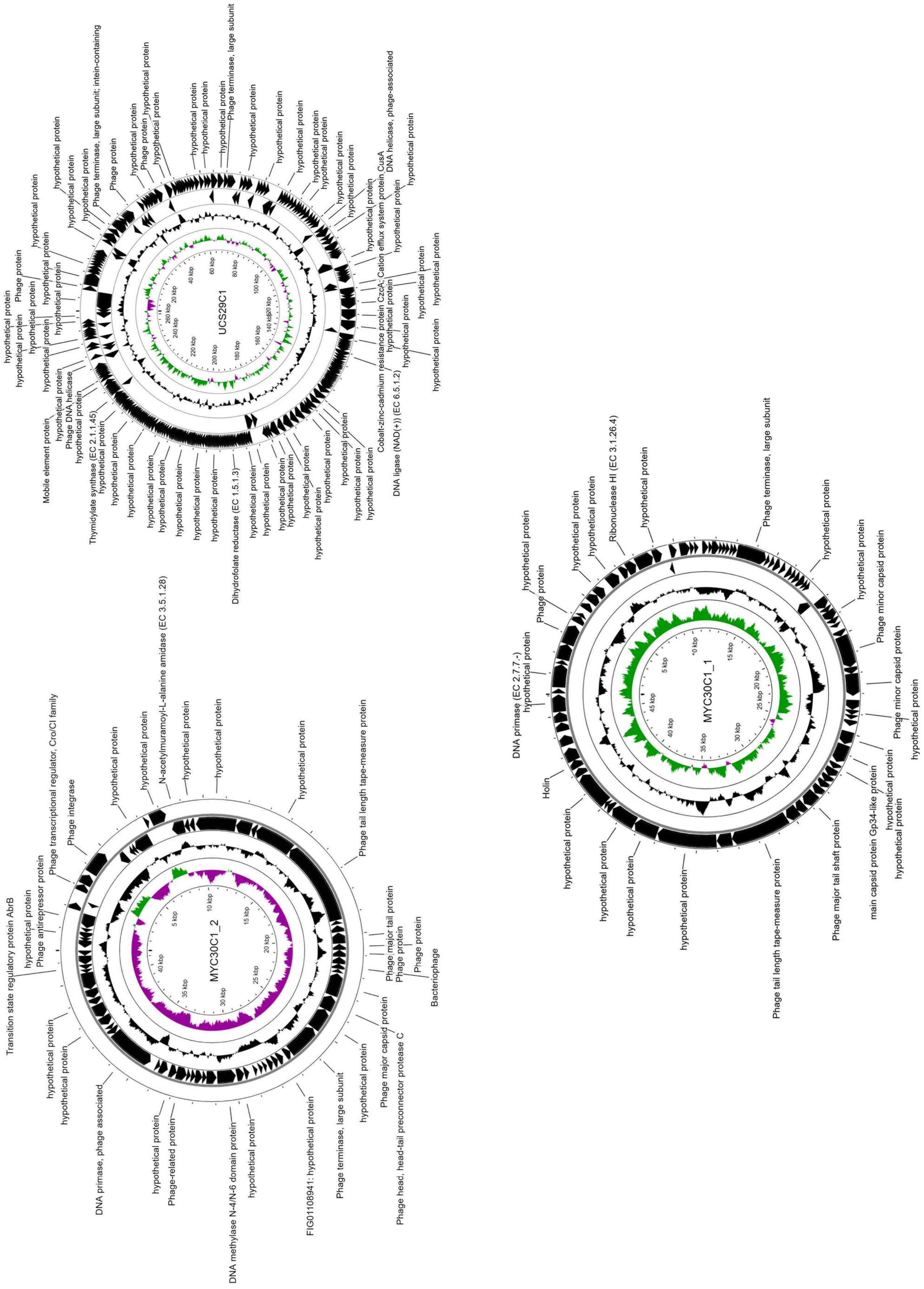


**Supplementary Figure 6: Annotated genomes of unique ΦHD-isolated phages UCS29C1, MYC30C1_1 and MYC30C1_2.** Organization of UCS29C1, MYC30C1_1 and MYC30C1_2 genomes. Circular genome map was displayed with CGView. Genomes were annotated using **RAStk.** Coding sequences (CDS, black) is depicted on the two outer rings. **GC content** is shown in black in the middle ring, while **GC skew** is represented on the two innermost rings, with positive skew in green and negative skew in purple.

**Supplemental Figure 7.**

**A.**

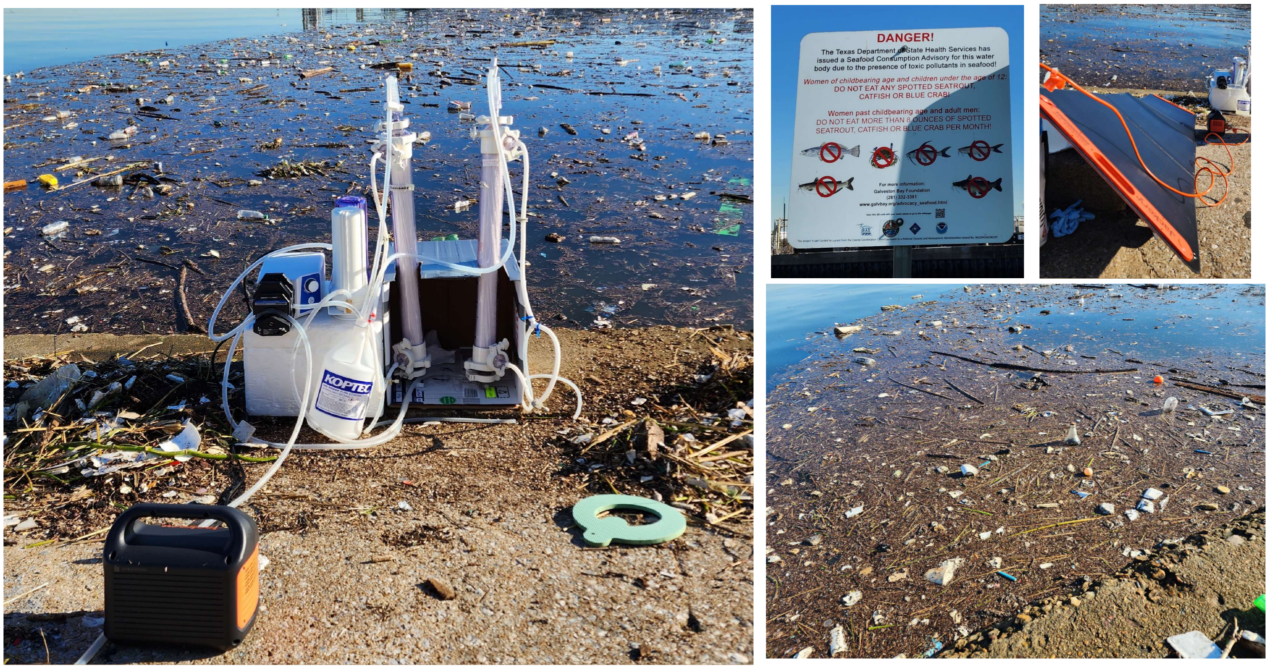


**B.**

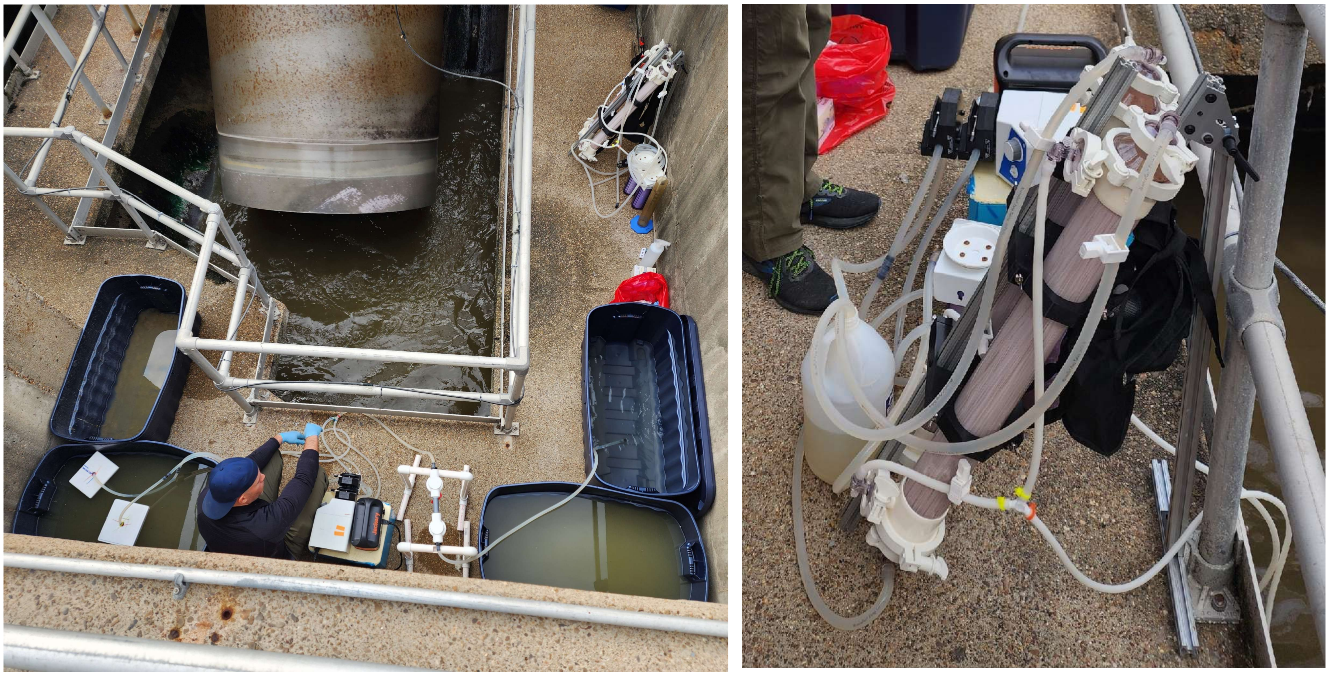

**Supplementary Figure 7: Field photos of ΦHD**. **A)** Buffalo Bayou is accessible at Santa Ana Capture Site. It is relatively close to the Port of Houston where ships enter and exit the Gulf of Mexico. Generally, this location is polluted with trash and dead plant material. There is also a warning sign that cautions women and children to not eat any spotted seatrout, catfish, or blue crab caught in the Bayou due to toxic pollutants present. **B)** West University WWTP (influent). Raw wastewater enters the plant through large pipes where they are subjected to a large filter (steel bars) to filter hygiene products and trash. In the picture, the influent collects in this container and is propelled upwards with a large Archimede's screw (towards the contact tank).

### Supplementary Tables 1-3

| Bacterial Isolates | Distinct Count of Case Name | Number of Isolates |
|---|---|---|
| *Achromobacter xylosozidans* | 6 | 10 |
| *Acinetobacter baumanii* | 3 | 5 |
| *Burkholderia gladioli* | 1 | 1 |
| *Burkholderia spp* | 1 | 1 |
| *Enterobacter cloacae* | 4 | 7 |
| *Enterococcus faecalis* | 2 | 2 |
| *Enterococcus spp* | 6 | 9 |
| *Escherichia spp* | 3 | 5 |
| *Escherichia coli* | 57 | 92 |
| *Klebsiella aerogenes* | 3 | 3 |
| *Klebsiella oxytoca* | 1 | 2 |
| *Klebsiella pneumoniae* | 41 | 71 |
| *Morganella morganii* | 1 | 2 |
| *Proteus mirabilis* | 1 | 1 |
| *Providencia stuartii* | 1 | 1 |
| *Pseudomonas aeruginosa* | 74 | 153 |
| *Raotulla ornitholytica* | 1 | 2 |
| *Serratia marcescens* | 3 | 3 |
| *Staphylococcus aureus* | 29 | 53 |
| *Staphylococcus epidermidis* | 8 | 11 |
| *Staphylococcus lugdensis* | 1 | 3 |
| *Staphylococcus warneri* | 1 | 1 |
| *Stenotrophomonas maltophilia* | 8 | 15 |
| *Streptococcus intermedius* | 1 | 1 |
| *Streptococcus mitis* | 1 | 1 |
| *Unknown* | 5 | 7 |
| **Grand Total** | **263** | **462** |

**Supplementary Table 1.** TAILOR distinct case count and number of isolates in their bacterial library

| Bacterial Target | Count of Phages from the TAILOR Library |
|---|---|
| *Achromobacter xylosozidans* | 22 |
| *Acinetobacter baumanii* | 4 |
| *Enterobacter cloacae* | 9 |
| *Entercoccus faecalis* | 10 |
| *Enterococcus spp* | 5 |
| *Escherichia coli* | 64 |
| *Klebsiella aerogenes* | 4 |
| *Klebsiella pneumoniae* | 80 |
| *Pseudomonas aeruginosa* | 62 |
| *Raotulla ornitholytica* | 1 |
| *Serratia marcescens* | 4 |
| *Staphylococcus aureus* | 13 |
| *Staphylococcus pseudintermedius* | 20 |
| *Stenotrophomonas maltophilia* | 27 |
| **Grand Total** | **325** |

**Supplementary Table 2.** TAILOR phage library

| Bacterial Isolates | Number of isolates with NO phage |
|---|---|
| *Achromobacter xylosozidans* | 3 |
| *Acinetobacter baumanii* | 7 |
| *Enterobacter cloacae* | 2 |
| *Entercoccus faecium* | 5 |
| *Entercoccus spp* | 1 |
| *Escherichia coli* | 11 |
| *Klebsiella aerogenes* | 1 |
| *Klebsiella pneumonia* | 8 |
| *Proteus mirabillis* | 1 |
| *Providencia stuartii* | 1 |
| *Pseudomonas aeruginosa* | 35 |
| *Serratia marcescens* | 2 |
| *Staphylococcus aureus* | 10 |
| *Staphylococcus epidermidis* | 7 |
| *Staphylococcus lugdensis* | 3 |
| *Staphylococcus warneri* | 1 |
| *Stenotrophomonas maltophilia* | 2 |
| *Streptococcus intermedius* | 1 |
| **Grand Total** | **101** |

**Supplementary Table 3.** Number of isolates with NO phage in TAILOR's library

### Supplementary Table 4

| *Reference* | *Source* | *Sampling Strategy* | *Approximate Vol. Processed (Liters)* | *Total viral contigs or viral populations reported* |
|---|---|---|---|---|
| *Roux et al 2016. Nature* | Tara Oceans expedition<br>Malaspina (seawater) | 90 samples (20L each)<br>14 samples (80L each) | 2920L | 15,222 viral population (≥10kb) |
| *Gios et al 2024* | Groundwater from two aquifers in Canterbury, New Zealand, | 59 wells (3 to 90L groundwater per well) | 177-5310L | 5,672 (≥5kb), 468 (≥10 kb) vOTUs |
| *Xiong et al 2023* | Freshwater from Napahai plateau wetland | 8 sampling points (25kg/site)+soil | 200L | 5,129 (≥1 kb), 199 (≥10kb) vOTUs |
| *Chen et al. 2023* | Freshwater from Middle Route of the South-to-North Water Diversion Canal, China | 30L/sample (64 total) | 1920L | 40,261 (≥5 kb) vOTUs |
| *Labbe et al 2020* | Fresh/Seawater from Lakes on Ellesmere Island in the Canadian High Arctic | 375-900 mL per sample (5 samples, triplicates) | ~20L | 16,080 (≥2kb) vOTUs |
| *Angly et al. 2006* | Seawater from Sargasso Sea, Gulf of Mexico, British Columbia, and Artic Ocean | Sargosso Sea (SAR): 1 sample (150L)<br>Gulf of Mexico (GOM): 41 samples (20-100L per sample)<br>Bay of British Colombia (BBC): 85 samples (20-100L per sample)<br>Arctic Ocean: 56 samples (20-100L per sample) | ~2,000-37,000L | 251,098 "sequences" identified as viral (14.2% of 1768297 total sequences identified as "viral") |
| *Malki K. et al 2020* | Ichetucknee Springs, Jackson Springs, Manatee Springs, Rainbow Springs, and Volusia Springs (Florida, USA) | 50L per site in triplicates | ~750L | 55,254 viral contigs (≥1kb) |
| *Schoenfeld et al. 2008* | Bear Paw and Octopus Hot Springs, California, USA | . | 400-600L | 2710 "viral types" |
| *Liu et al. 2024* | Wastewater (WWTPs, Texas, USA) | Collection of influent and anaerobic digestion from various sites (300-500 mL per site) | 16L | 1322 viral contigs (≥5kb) |
| | Freshwater (Texas, USA) | 57L at 4 sites | 220L | 2780 viral contigs (≥5kb) |
| | Seawater (Texas, USA) | 57L at 2 sites | 110L | 9676 viral contigs (≥5kb) |
| *This study* | Wastewater from West Uni. WWTP | . | 400L | 16,198 (≥5kb), 5095 (≥10kb) vOTUs |
| | Freshwater from Brays Bayou | . | 400L | |
| | Freshwater from Clear Creek | . | 400L | |
| | Seawater from West Bay, Galveston, USA | . | 400L | |

**Supplementary Table 4: Various viral metagenomic studies with approximate volume processed and reported viral contigs.** A total of 16,198 virus operational taxonomic units (vOTUs) were identified from four specific sites in this studies' metagenomic datasets. 5,095 vOTUs were ≥10 kb. Viral contigs were identified from trimmed and cleaned reads with VIBRANT. vOTUs were clustered with standard thresholds of 95% average nucleotide identity over 85% alignment fraction. Additional selected references are shown to depict total volume sampled, viral contigs/viral populations present, and sampling strategy used.

### Supplementary Table 5-8

| | Date Processed | Volume | Concentration | Titer (PFU/mL) | vOTUs (≥ 5kb / ≥ 10kb) |
|---|---|---|---|---|---|
| **West Uni. WWTP** | 10/16/2024 | 400L | 2000X | 1.03E+06 | 3355 / 881 |
| **Brays Bayou (H.E.B)** | 8/22/2024 | 400L | 4000X | 2.59E+04 | 4173 / 1541 |
| **Clear Creek (Challenger Park)** | 8/23/2024 | 400L | 4000X | 4.33E+03 | 3898 / 1259 |
| **West Bay (Galveston)** | 10/1/2024 | 400L | 5333X | / | 4772 / 1414 |
| **Hamilton Pool Preserve** | 1/21/2023 | 305L | 4363X | 1.10E+02 | 3822 / 1604 |
| **Buffalo Bayou** | 3/8/2023 | 600L | 22222X | 6.19E+04 | 3594 / 1262 |
| **Pedernales State Park** | 1/20/2023 | 774L | 15486X | 1.14E+03 | 1361 / 3336 |
| **Brays Bayou (MacGregor Park)** | 6/2/2023 | 328L | 6560X | / | 4601 / 1782 |
| **Clear Creek (Walter Hall Park)** | 6/2/2023 | 364L | 4853X | / | 3815 / 1328 |
| **Brays Bayou (HEB) 2nd** | 8/27/2023 | 400L | 5333X | / | 16418 / 6156 |

**Supplementary Table 5: Environmental sampling summary.** Water from various sites have been processed with ΦHD and further concentrated in-lab; volume processed and concentration after second stage processing are listed. Concentrates were titered on indicator strain *P. aeruginosa* PA01. DNA was extracted from 11.5 mL of each concentrate and sent for PCR-free shallow shotgun metagenomic sequencing. After metagenomic analysis, viral contigs were identified and clustered into vOTUs based on 5kb and 10kb cut-offs. Raw sequences were uploaded to SRA (BioProject #: PRJNA1308632).

| | Metapop derived α-diversity calculations | | | | | | | | β-diversity |
|---|---|---|---|---|---|---|---|---|---|
| | Richness | Shannons_H | Simpson | InvSimpson | Fisher | Pielous_J | Chao1 | ACE | Brays-Curtis Dissimilarity |
| 5L | 21918 | 8.8771 | 0.9988 | 826.6507 | 4172.3304 | 0.8882 | 21918.5969 | 21915.5849 | 0.6062 |
| 400L | 21918 | 8.7726 | 0.9986 | 690.5897 | 4154.8586 | 0.8777 | 21918.1875 | 21918.3995 | |

**Supplementary Table 6: α-diversity analysis for metagenomic datasets of unprocessed (5L sample) and processed** ϕHD concentrate **(60 mL 6667X concentrate) from freshwater.** We used MetaPop's macrodiversity pipeline to calculate diversity.

| Source | Total Volume Processed | Concentrates after 2nd stage processing | vOTUs ≥ 5kb / ≥ 10kb | Number of µg sent for sequencing |
|---|---|---|---|---|
| **Freshwater from Brays Bayou** | 400L | 6667X | 16418 / 6156 | 184.4 |
| **Freshwater from Brays Bayou** | 5L | / | 5500 / 2089 | 25.6 |

**Supplementary Table 7: Viral counts of unprocessed vs. processed concentrates.** We compared the viromes of unprocessed and ΦHD processed freshwater samples. ΦHD was used to process 400L of freshwater from Brays Bayou and further concentrated in-lab. Final concentration was 6667X, and the total volume was used for DNA extraction. 5L of unprocessed freshwater was concentrated down to 60 mL and used for DNA extraction. DNA was sent for PCR-free shallow shotgun metagenomic sequencing.

| | Metapop derived α-diversity calculations | | | | | | | |
|---|---|---|---|---|---|---|---|---|
| | Richness | Shannons_H | Simpson | InvSimpson | Fisher | Pielous_J | Chao1 | ACE |
| Freshwater (Brays Bayou) | 6069 | 7.6981 | 0.9986 | 719.9960 | 941.3756 | 0.8837 | 5006.4854 | 4860.2307 |
| Brackish (Clear Creek) | 5788 | 7.6350 | 0.9983 | 575.8816 | 895.3257 | 0.8813 | 4815.2402 | 4626.3387 |
| Seawater (Galveston) | 5172 | 7.5107 | 0.9971 | 348.8557 | 923.8425 | 0.8783 | 4887.0294 | 4855.5378 |
| Wastewater (West University WWTP) | 3721 | 7.7792 | 0.9993 | 1372.8294 | 780.3868 | 0.9462 | 3506.4615 | 3466.4603 |

**Supplementary Table 8: α-diversity analysis for metagenomic datasets of brackish, sea, waste, and freshwater**. We used MetaPop's macrodiversity pipeline to calculate diversity.

### Supplementary Table 9

| Phage | Accession Number | Coverage | length (bp) | Nearest neighbour, origin and source (if available) | Query Coverage | Percent Identity | tANI | Neighbor Accession Number | Comments |
|---|---|---|---|---|---|---|---|---|---|
| **E coli** | | | | | | | | | |
| UCS29C1 | PZ285958 | 2758 | 261881 | Phage Miami, Houston, TX, sewage | 85 | 95 | 81 | NC_070950.1 | new species |
| UCS29C2 | PZ285959 | 12363 | 60722 | Klebsiella Phage PIN1, , waste water Australia | 84 | 95 | 80 | PQ803402.1 | new species |
| UCS37C1 | PZ285960 | 11376 | 10260 | vB_EcoM_DE17, waste water from China | 99 | 99 | 98 | OP595146.1 | incomplete genome |
| UCS37C2_1 | PZ229304 | 112 | 149154 | vB_Eco_J-01, Monterrey, Mexico, CF, intestine | 96 | 99 | 95 | PQ438393.1 | new strain |
| UCS37C2_2 | PZ285961 | 11839 | 17642 | vB_EcoM_DE17 and UCS37C1 above | 99 | 100 | 99 | same as UCS37C1 | |
| BSL02C1_1 | PZ285931 | 1216 | 165758 | JLBYU31, Provo, Utah | 97 | 97 | 94 | OK272484.1 | new species |
| BSL02C1_2 | PZ285932 | 7547 | 25101 | vB_Eco-DE17, and UCS37C1 above | 99 | 98 | 97 | same as UCS37C1 | |
| **Enterococcus** | | | | | | | | | |
| MYC30C1_1 | PZ285937 | 18138 | 46030 | Bacillus phage PBC1 (1%), Seoul, Korea | 1 | 77 | 1 | NC_017976.1 | new genus |
| MYC30C1_2 | PZ285938 | 19 | 41467 | Bacillus phage SDFMU_Pfc (43%), China, Housefly intestines | 38 | 94 | 36 | OQ884029.1 | new genus |
| UCS17X2C1 | PZ285939 | 10822 | 56145 | vB_OCTPT_PG2, 95%, UC Irvine, sewage | 95 | 96 | 91 | ON113177.1 | new species |
| UCS17X2C2 | PZ285940 | 8161 | 56896 | Enterococcus phage vB_Efas_TV16, Rome, sewage | 94 | 97 | 91 | NC_041959.1 | new species |
| **Klebsiella** | | | | | | | | | |
| UCS26C1_1 | PZ285948 | 8458 | 48577 | vB_Kpn_K28PH129, Spain, wastewater | 99 | 97 | 96 | OY757089.1 | new strain |
| UCS26C1_2 | PZ285949 | 3973 | 43302 | Klebsiella phage Kpn_BHU3, River Ganga, India | 90 | 95 | 86 | OL976437.1 | new species |
| EUA02C1 | PZ285971 | 329 | 41069 | Klebsiella phage P79_1, China, wastewater | 91 | 94 | 86 | OR256027.1 | new species |
| EUA3C1 | PZ285966 | 5556 | 40104 | aligns well to EUA02C1 | NA | NA | N/A | NA | same as EUA02C1 |
| MYC16C2 | PZ285936 | 8733 | 59447 | Klebsiella phage RCIP0070, China, wastewater | 88 | 97 | 85 | OR532864 | new species |
| UCS26X1C1_1 | PZ285952 | 8839 | 47157 | phi731, Hungary | NA | NA | N/A | NA | same as UCS26C1_1 |
| UCS26X1C1_2 | PZ285972 | 11054 | 27674 | phi1_146045, China, sewage, probably the same as UCS26C1_2 | NA | NA | N/A | NA | same as UCS26C1_2 |
| UCS26X1C2_1 | PZ285953 | 1423 | 49347 | vB_Kpn_K28PH129, Spain | NA | NA | N/A | NA | same as UCS26C1_1 |
| UCS26X1C2_2 | PZ285954 | 416 | 24431 | CPRSB, Kenya, sewage | 99 | 98 | 97 | OM971649.1 | incomplete genome |
| UCS26X1C2_3 | PZ285955 | 402 | 35324 | vB_KpnM_Satellite_ER45, Chicago, sewage | 98 | 99 | 97 | PP738781.1 | new strain |
| UCS26X1C2_4 | PZ285956 | 401 | 19900 | CPRSA, Kenya, sewage | 95 | 97 | 92 | OM971648 | incomplete genome |
| UCS26X1C2_5 | PZ285970 | 26306 | 8196 | vB_KpnS_Carter_MM5, Chicago, sewage | 99 | 98 | 97 | PP738797.1 | same as USC26C2_1 |
| UCS26X1C2_6 | PZ285957 | 22041 | 8074 | phiKp_14, Japan, sewage | 98 | 97 | 95 | LC768476.1 | same as USC26C2_1 |
| UCS26C2_1 | PZ285950 | 13393 | 46145 | vB_KpnS_mingus_MM1, Chicago, sewage | 96 | 97 | 93 | PP738793.1 | new strain |
| UCS26C2_2 | PZ285951 | 5964 | 49413 | KpKT21Phi1, Israel | 97 | 96 | 93 | NC_048143.1 | new species |
| UCS26C2_3 | PZ285973 | 2594 | 27370 | phi1_146045, China, Sewage, probably identical to UCS26C1_2 | NA | NA | N/A | NA | same as UCS26C1_2 |
| MYC16C1 | PZ285935 | 11903 | 43492 | VB_KM5a1-JVSB2, Ohio, soil | 95 | 96 | 91 | PP444685.1 | new species |
| **Pseudomonas** | | | | | | | | | |
| CSM01C1 | PZ285933 | 356 | 10834 | PSA11, Pittsburg, sewage | 95 | 99 | 94 | MZ08973.1 | incomplete genome |
| UCS20C3 | PZ285945 | 357 | 26742 | vB_PaeP_130_113, Naval Research Center, Fort Detrick, MD,USA | 99 | 95 | 94 | NC_047953.1 | incomplete genome |
| UCS18C3_1 | PZ285941 | 708 | 12737 | HZ2201, China | 100 | 97 | 97 | OQ427617.1 | incomplete genome |
| UCS18C3_2 | PZ285942 | 1535 | 2060 | PaGz-1, China, freshwater | 100 | 100 | 100 | NC_073603.1 | incomplete genome |
| UCS20C5 | PZ285947 | 216 | 44853 | Epa1, Walter Reed, USA | 99 | 97 | 96 | MT108723.1 | new strain |
| UCS20C4 | PZ285946 | 7654 | 3918 | WP1, West Point, USA | 100 | 91 | 91 | PP596839.1 | incomplete genome |
| 6X1C4 | PZ285969 | 4137 | 179890 | new genus, Astolliot, Denmark, lake/soil, 4% coverage | 4 | 81 | 3 | OM982621.1 | new genus |
| UCS20C1 | PZ285944 | 235 | 45086 | ZCPS1, Egypt, sewage | 99 | 98 | 97 | ON156559.1 | new strain |
| MGB1X4C4 | PZ285934 | 1352 | 19227 | WP1, West Point, USA | 99 | 90 | 89 | PP596839.1 | incomplete genome |
| MGB1X4C1 | PZ285968 | 16890 | 40393 | debbie, Denmark, sewage | 97 | 98 | 95 | MT119363.1 | new strain |
| UCS18C4 | PZ285943 | 12336 | 84317 | vB_PaeM_VL12, Thailand, pond | 97 | 98 | 95 | OM421595.1 | new strain |
| MGB1X4C3 | PZ285967 | 13500 | 65863 | WP1, West Point, USA, | 99 | 92 | 91 | PP596839.1 | new species |

**Supplemental Table 9: Characteristics of sequences assembled from members of the R-Phage library.** Nearest neighbors determined by total score (BLASTn), coverage and identity also from BLASTn output. Determination of assemblies representing the same phage by nucleotide alignment (BLASTn), proteomic clustering (VIPTree) plaque morphologies and known origins of the host. Phages highlighted with the same color indicate they are identical. Submission of whole genome sequences to GenBank in progress.

**Supplementary Table 11**

<table>
<tr><td colspan="7">Sampling for GEΦMAPPING</td></tr>
<tr><td rowspan="2">Water</td><td rowspan="2">Location</td><td rowspan="2">Latitude</td><td rowspan="2">Longitude</td><td colspan="3">Date Collected in MM/DD/YYYY</td></tr>
<tr><td colspan="3">Temperature(High°F/Low°F); Precipitation (in)</td></tr>
<tr><td>Freshwater</td><td>Sims Bayou @ Townwood Park</td><td>29.619</td><td>-95.43</td><td rowspan="14">10/24/2023 (56/48); 0.0</td><td rowspan="14">8/30/2024 (87/76); 0.48</td><td rowspan="15">9/19/2024 (95/78); 0.0</td></tr>
<tr><td>Brackish water</td><td>Clear Creek @ Walter</td><td>29.514</td><td>-95.104</td></tr>
<tr><td>Seawater</td><td>Gulf of Mexico, Galveston</td><td>29.267</td><td>-94.823</td></tr>
<tr><td>Freshwater</td><td>Vince Bayou @ Pasadena Fire Station 4</td><td>29.677</td><td>-95.21</td></tr>
<tr><td>Brackish water</td><td>Buffalo Bayou @ Santa Ana Capture Site</td><td>29.725</td><td>-95.212</td></tr>
<tr><td>Seawater</td><td>West Bay, Galveston</td><td>29.289</td><td>-94.874</td></tr>
<tr><td>Freshwater</td><td>Brazos River</td><td>29.375</td><td>-95.576</td></tr>
<tr><td>Freshwater</td><td>Buffalo Bayou @ Memorial Park</td><td>29.76</td><td>-95.439</td></tr>
<tr><td>Freshwater</td><td>White Oak Bayou @ TC Jester</td><td>29.828</td><td>-95.456</td></tr>
<tr><td>Freshwater</td><td>Halls Bayou @ Keith Wiess Park</td><td>29.895</td><td>-95.355</td></tr>
<tr><td>Freshwater</td><td>Green's Bayou @ Wtidwell</td><td>29.85</td><td>-95.228</td></tr>
<tr><td>Freshwater</td><td>Brays Bayou @ McGregor Park</td><td>29.714</td><td>-95.339</td></tr>
<tr><td>Freshwater</td><td>Brays Bayou @ Medical Center</td><td>29.708</td><td>-95.391</td></tr>
<tr><td>Brackish water</td><td>Clear Creek @ Challengar</td><td>29.506</td><td>-95.133</td></tr>
<tr><td>Freshwater</td><td>Bray's Bayou @ HEB</td><td>29.712</td><td>-95.379</td><td></td><td></td></tr>
<tr><td>Wastewater</td><td>Almeda Sims WTTP</td><td>29.629</td><td>-95.408</td><td rowspan="7">1/30/2024 (73/47); 0.0</td><td rowspan="7">3/1/2024 (73/53); 0.0</td><td rowspan="7">9/24/2024 (92/78); 0.0</td></tr>
<tr><td>Wastewater</td><td>Homestead WWTP</td><td>29.808</td><td>-95.295</td></tr>
<tr><td>Wastewater</td><td>International Airport Houston WWTP</td><td>29.965</td><td>-95.354</td></tr>
<tr><td>Wastewater</td><td>Metro Central WWTP</td><td>29.591</td><td>-95.155</td></tr>
<tr><td>Wastewater</td><td>Park Ten WWTP</td><td>29.79</td><td>-95.67</td></tr>
<tr><td>Wastewater</td><td>Turkey Creek WWTP</td><td>29.768</td><td>-95.622</td></tr>
<tr><td>Wastewater</td><td>West University WWTP</td><td>29.696</td><td>-95.42</td></tr>
<tr><td>Wastewater</td><td>Influent Tank, West Univeristy WWTP</td><td>29.696</td><td>-95.42</td><td rowspan="3">9/24/2024 92/78); 0.0</td><td rowspan="3" colspan="2"></td></tr>
<tr><td>Wastewater</td><td>Aeration Tank, West Univeristy WWTP</td><td>29.696</td><td>-95.42</td></tr>
<tr><td>Wastewater</td><td>Contact Tank, West Univeristy WWTP</td><td>29.696</td><td>-95.42</td></tr>
</table>

**Supplementary Table 10: Samples collected for GEOMAPPING from environmental and wastewater.** Samples collected for 16S sequencing were collected on the days recorded MM/DD/YYYY. Latitude/Longitude were record as well as temperature (High °F/ Low °F) and precipitation (inches). Raw sequences were uploaded to SRA (BioProject #: PRJNA1309115).

### Supplementary Table 11-12

| Short-hand | Strain | Characteristic | Reference |
|---|---|---|---|
| 17978 | *Acinetobacter baumanii 17978* | Isolated from fatal meningitis | ATCC (Bouvet and Grimont et al., 1987) |
| UTS1 | *Achromobacter xylosozidans UTS1* | Isolated cystic fibrosis (CF) lung infection | TAILΦR LABS (Terwilliger et al., 2020) |
| SLC1 | *Enterobacter cloacae SLC1* | Isolated from hip prosthesis wound | TAILΦR LABS (Terwilliger et al., 2020) |
| VRE001 | *Enterococcus faecalis VRE001* | Vancomycin-resistant isolate from patient stool | Haddad et al., 2019 |
| K12 | *Escherichia coli K12 (MG1655)* | Wild-type, common laboratory strain | |
| JJ2528 | *Escherichia coli JJ2528* | ExPEC ST131 H30-R | Price et al., 2013 |
| UCF1 | *Klebsiella aerogenes UCF1* | Isolated from hip prosthesis wound | TAILΦR LABS (Terwilliger et al., 2020) |
| EUA2 | *Klebsiella pneumonia EUA2* | Isolated from urinary tract infection (UTI) | TAILΦR LABS (Terwilliger et al., 2020) |
| DSA97 | *Pseudomonas aeruginosa DS497* | Isolate from UTI | Robert Frank Ramig |
| PA01 | *Pseudomonas aeruginosa PA01* | Common laboratory strain; Isolated from wound | Holloway, 1955 |
| AZM28 | *Staphylococcus aureus AZM28* | Isolate from dog skin swab #13-1 on MSA | TAILΦR LABS (Terwilliger et al., 2020) |
| AZM69 | *Staphylococcus pseudointermedius AZM69* | Coagulase positive; isolated from canine ear infection | TAILΦR LABS (Terwilliger et al., 2020) |
| AR0246 | *Pseudomonas aeurignosa AR0246* | MDR strain, CDC and FDA Antibiotic Resistance Isolate Bank | TAILΦR LABS (Terwilliger et al., 2020) |
| AR0266 | *Pseudomonas aeurignosa AR0266* | MDR strain, CDC and FDA Antibiotic Resistance Isolate Bank | TAILΦR LABS (Terwilliger et al., 2020) |
| BSL02 | *Escherichia coli BSL01* | Isolated from UTI | TAILΦR LABS |
| UCS29 | *Escherichia coli UCS29* | Isolated from UTI | TAILΦR LABS |
| UCS37 | *ESBL Escherichia coli UCS37* | Isolated from urine (pyelonephritis, sepsis) | TAILΦR LABS |
| UCF06 | *ESBL Escherichia coli UCF06* | Isolated from bone (infected amputation) | TAILΦR LABS |
| MYC16 | *Klebsiella aerogenes MYC16* | Isolated from shoulder prosthetic joint infection | TAILΦR LABS |
| EUA02 | *Klebsiella pneumoniae EUA01* | Isolated from UTI | TAILΦR LABS |
| EUA03 | *Klebsiella pneumoniae EUA02* | Isolated from UTI | TAILΦR LABS |
| UCS26 | *Klebsiella pneumoniae UCS26* | Isolated from UTI, bacteremia | TAILΦR LABS |
| UCS26.1 | *Klebsiella pneumoniae UCS26.1* | Same as UCS26 | TAILΦR LABS |
| MYC20 | *MSSA Staphylococcus aureus MYC20* | Isolated from sputum (bronchiectasis) | TAILΦR LABS |
| UCS17.2 | *Escherichia coli UCS17.2* | Isolated from UTI | TAILΦR LABS |
| CSM01 | *Pseudomonas aeruginosa CSM01* | Isolated from sputum (lung abscess, adjacent sternal osteomyelitis) | TAILΦR LABS |
| UCS18 | *Pseudomonas aeruginosa UCS18* | Isolated from bile duct (biliary infection, sepsis) | TAILΦR LABS |
| UCS20 | *Pseudomonas aeruginosa UCS20* | Isolated from chest drain (lung, chest wall infection) | TAILΦR LABS |
| UCS23 | *Pseudomonas aeruginosa UCS23* | Isolated from UTI | TAILΦR LABS |
| MGB1.4 | *Pseudomonas aeruginosa MGB1.4* | Isolated from sputum (CF lung) | TAILΦR LABS |
| HPC3.1 | *Pseudomonas aeruginosa HPC3.1* | Isolated from sputum (lung) | TAILΦR LABS |
| 17802 | *Vibrio parahaemolyticus* 17802 | Isolated from Shirasu food poisoning | ATCC (Fujini et al., 1974) |
| 27562 | *Vibrio vulnificus* 27562 | Isolated from human blood | ATCC (Reichet et al., 1976) |

**Supplementary Table 11: Strains used in the study.** We include short-hand names, full strain name, characteristics, and references for the strains.

| Phage | Accession Number | Coverage | Length (bp) | Nearest neighbour, origin and source (if available) | Query coverage | Percent Identity | tANI | Neighbor Accession Number | Comments |
|---|---|---|---|---|---|---|---|---|---|
| VP1_GalvWB | PZ285965 | 12922 | 42735 | VP48, China, seawater | 87 | 77 | 66.99 | KX371615.1 | new species |
| VP2 | PZ285962 | 26940 | 43347 | CAUVibPm, Sorae Beach, South Korea, seawater | 100 | 92.94 | 92.94 | PZ038745 | new species |
| VP3 | PZ285963 | 26421 | 42657 | VP48, China, seawater | 87 | 77 | 66.99 | KX371615.1 | likely same as VP1 |
| VP4 | PZ285964 | 24062 | 42812 | VvB_VpP_AC2, China, seawater | 99 | 93.45 | 92.5155 | ON864052.1 | very similar (98%) to VP2, likely different strain |

**Supplementary Table 12: Additional phages found from this study.** Nearest neighbors determined by total score (BLASTn), coverage and identity also from BLASTn output. Determination of assemblies representing the same phage by nucleotide alignment (BLASTn), proteomic clustering (VIPTree) plaque morphologies and known origins of the host. Phages highlighted with the same color indicate they are identical. Submission of whole genome sequences to GenBank in progress
